# Biased cell adhesion organizes a circuit for visual motion integration

**DOI:** 10.1101/2023.12.11.571076

**Authors:** Yannick Carrier, Laura Quintana Rio, Nadia Formicola, Vicente de Sousa-Xavier, Maha Tabet, Yu-Chieh David Chen, Maëva Wislez, Lisa Orts, Filipe Pinto-Teixeira

**Affiliations:** MCD, Centre de Biologie Intégrative (CBI), CNRS, Université de Toulouse, UT3, Toulouse, France; Department of Biology, New York University, New York, NY 10003, USA

## Abstract

Layer specific computations in the brain rely on neuronal processes establishing synaptic connections with specific partners in distinct laminae. In the *Drosophila* lobula plate neuropile, the axons of the four subtypes of T4 and T5 visual motion direction-selective neurons segregate into four layers, based on their directional preference, and form synapses with distinct subsets of postsynaptic neurons. Four bi-stratified inhibitory lobula plate intrinsic cells exhibit a consistent synaptic pattern, receiving excitatory T4/T5 inputs in one layer, and conveying inhibitory signals to an adjacent layer. This layered arrangement establishes motion opponency. Here, we identify layer-specific expression of different receptor-ligand pairs belonging to the Beat and Side families of Cell Adhesion Molecules (CAMs) between T4/T5 neurons and their postsynaptic partners. Genetic analysis reveals that Beat/Side mediated interactions are required to restrict T4/T5 axonal innervation to a single layer. We propose that Beat/Side contribute to synaptic specificity by biasing adhesion between synaptic partners before synaptogenesis.

## INTRODUCTION

The clustering of synaptic connections into stereotyped layers is a hallmark of neural connectivity in complex nervous systems. The prevalence of layered synaptic arrangements in both invertebrates and vertebrates implies a fundamental organizational principle in neural development (Baier, 2013; Fischbach and Dittrich, 1989; Huberman et al., 2010; Morante and Desplan, 2008). However, the contribution of the layer assembly process to synaptic specificity remains unclear. Elucidating the mechanisms that govern the arrangement of neurons into layers, and their interplay with the mechanisms establishing synaptic specificity, is central to understanding how layer specific synaptic networks are built. To address this question, we focused on the development of the synaptic layered connectivity of the lobula plate neuropile of the *Drosophila* optic lobe. In the fly optic lobe, developmentally related T4 and T5 cells are the first visual motion direction-selective neurons (Apitz and Salecker, 2015; Oliva et al., 2014; Pinto-Teixeira et al., 2018). Both T4 and T5 neurons exist in four subtypes, each tuned to one of the four cardinal directions of visual motion. Neurons sharing the same directional tuning project their axons retinotopically to specific layers of the lobula plate. This arrangement results in four layers specialized for motion in each cardinal direction: two layers for horizontal motion (layer 1, front-to-back; layer 2, back-to-front) and two layers for vertical motion (layer 3, upward; layer 4, downward) (Maisak et al., 2013) (**Figure 1A**). A recent EM reconstruction of the lobula plate provided a comprehensive identification of neurons that receive input from T4/T5 neurons (Shinomiya et al., 2022). There are a total of 58 putative connected neuron types, 74% of which synapse with both T4 and T5 neurons of the same subtype (**Figure 1B**, **S1**). In each layer, a distinct set of downstream visual projection neurons (VPNs) supports the parallel processing of direction selective signals for a variety of roles in visually guided behaviors (Hausen, 1984; Klapoetke et al., 2017; Panser et al., 2016; Wu et al., 2016). For example, layer-specific Lobula Plate Columnar (LPC) and Lobula-Lobula Plate Columnar (LLPC) neurons establish a translational optic flow detection circuit downstream of T4/T5 neurons (Isaacson et al., 2023). Furthermore, in each layer, Lobula Plate Tangential Cells (LPTCs) integrate local motion signals into a global optic flow that matches the direction selectivity of their upstream T4/T5 neurons (Mauss et al., 2014; Schnell et al., 2012) (**Figure 1C, G**). Importantly, adjacent layers represent opposite directions of motion. This organization allows bi-stratified, inhibitory Lobula Plate Intrinsic (LPi) neurons, to receive input from T4/T5 in one layer, and convey inhibitory signals to LPTC cells in the neighboring motion-opponent layer (**Figure 1C-G**, **S2**) (Fischbach and Dittrich, 1989; Raghu and Borst, 2011). This layered organization establishes motion opponency and increases flow-field selectivity (Mauss et al., 2015). Thus, layers share a fundamental circuit pattern, consisting of T4/T5 neurons, a bilayer LPi cell, and several output VPNs.

**Figure 1.**
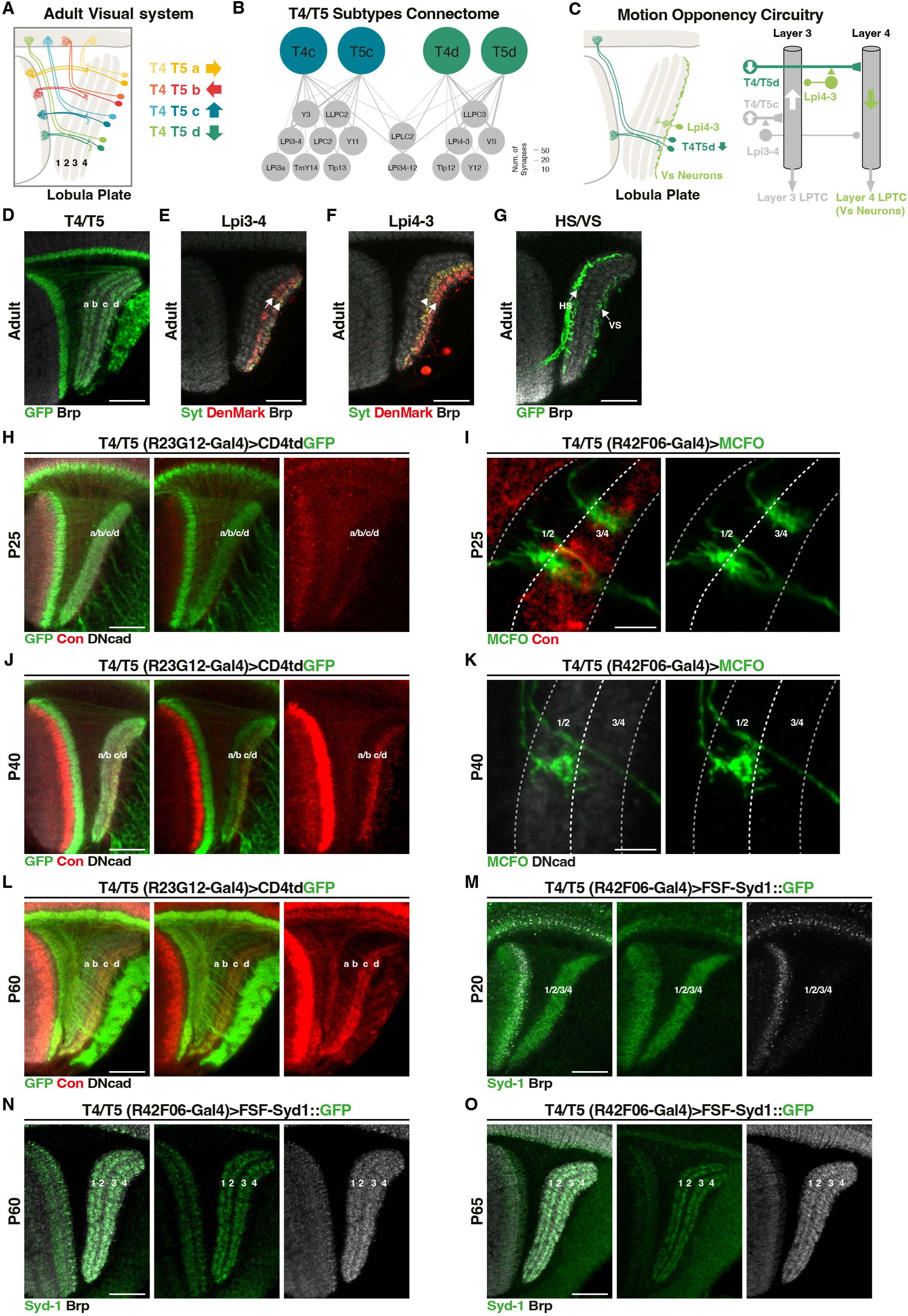
Lobula plate Motion Opponency Circuit. (**A**) Morphology of the four T4/T5 subtypes. All T4/T5 neurons with the same directional tuning preference project their axons retinotopically to the same layer of the lobula plate neuropile, producing four layers tuned for motion in each of the cardinal directions: layer 1, front-to-back; layer 2, back-to-front, layer 3, upward; layer 4, downward. (**B**) The connectome of T4/T5c and d subtypes with more than 10 synapses with a single T4 or T5 neurons, after (Shinomiya et al., 2022). (**C**) Motion Opponency Circuit in the lobula plate layer 4. Direction-selective wide-field motion responses in postsynaptic partners VS-Lobula Plate Tangential Cells (LPTCs) match the direction selectivity of their upstream T4/T5d neurons. LPi4-3 neurons receive input from T4/T5d neurons, and convey inhibitory signals to LPTC cells in the neighboring motion-opponent layer 3. LPi3-4 neurons contribute to a complementary circuit in the other direction (also, **Figure S2**). (**D**) T4/T5 neurons expressing membrane-GFP (green) (**E**) LPi3-4 neurons. (**F**) LPi4-3 neurons. (**G**) HS (layer 1) and VS (layer 4) LPTC neurons. In (**D-G**). Anti-Brp immunostaining (gray) labels the neuropiles. In (**E**) and (**F**) Syt::GFP labels axonal terminals (arrowhead) and the somatodendritic DenMARK::RFP labels the neurons, with stronger dendritic expression (arrow). (**H**-**L**) Sequential segregation of T4/T5 axons in four lobula plate layers during development. (**I**-**K**) **MCFO-**sparsely labeled T4/T5s during development. (**M-O**) In green, endogenous Syd1 expression in T4/T5 neurons during development through combination of a T4/T5 driver (R42F06-Gal4)>uas-flp and Frt-Stop-Frt-Syd1::GFP. In grey anti-Brp immunostaining. Scheme in (**C**) after (Mauss et al., 2015). In (I) and (K) grey dashed lines delineate the lobula plate, white dashed lines delineate a/b and c/d layers. Scale bars are 20μm, except in (I) and (K) where they are 5μm.

Using a published single cell RNAseq (scRNAseq) dataset of the developing optic lobe (Ozel et al., 2021), we found that axons of each T4/T5 subtype and their respective layer specific synaptic partners express matching binding partners of the Beat and Side proteins of the Immunoglobulin Superfamily (IgSF). Beat and Side proteins form a heterophilic protein interaction network (Li et al., 2017; Ozkan et al., 2013), whose founding members (Side and Beat-Ia) were identified as regulators of motor axon guidance in *Drosophila* embryos (Fambrough and Goodman, 1996; Siebert et al., 2009; Sink et al., 2001). The functions of the other paralogs remain unknown. Downregulation of these Beat/Side proteins in T4/T5 neurons results in the loss of layer-specific targeting – T4/T5 axons target their default layer, and also the adjacent layer that normally they do not innervate. Conversely, the ectopic expression of Beat/Side in the T4/T5 LPi postsynaptic neurons is sufficient to change LPi layer innervation, leading to the development of ectopic synapses with T4/T5 neurons in different layers. Crucially, synchronized Beat/Side expression between synaptic partners sustains T4/T5 layer segregation, as decreasing layer-specific Beat/Side in T4/T5 or in all postsynaptic neurons disrupts T4/T5 single layered innervation. Our findings support a model in which Beat/Side are not required for synapse formation, but rather contribute to synaptic specificity by biasing T4/T5 axons adhesion to layer specific synaptic partners prior to synaptogenesis. We further propose that, in each layer, the emergent synaptic specificity of each neuron depends on a competitive dynamic involving shared Beat/Side expression among all postsynaptic partners, each striving to establish connections with T4/T5 neurons.

## RESULTS

### Synchronous innervation of T4/T5 neurons and their postsynaptic partners during lobula plate development

T4 and T5 neurons are the most numerous cell types innervating the lobula late, and serve as the primary presynaptic neurons in all four lobula plate layers (**Figure 1A**). Given their number and functional significance, we postulated that T4/T5 neurons play a pivotal role in the structural foundation of lobula plate development. To test this hypothesis, we eliminated all T4/T5 neurons by driving expression of the pro-apoptotic gene hid using the R23G12-Gal4 driver (Kurmangaliyev et al., 2019), which is exclusively expressed in all T4/T5 progenitors and T4/T5 neurons from the third instar larval stage (L3) (**Figure S3**). We found that in the absence of T4/T5 neurons, the lobula plate is not formed (**Figure S3**), confirming a essential role of T4/T5 neurons in the development of this neuropile. We then set out to determine the timing of layer formation and the establishment of innervation by T4/T5 neurons and their postsynaptic partners, along with the restriction of synaptic connections to specific layers during development. Our analysis starts post-neurogenesis, at 25% pupation (P25). First, we visualized T4/T5 axonal projections using R23G12-Gal>CD4tdGFP and co-stained against the CAM Connectin (Con), which specifically labels the vertical motion sensing neurons layers 3 and 4 (Gao et al., 2008). At this developmental stage, the axonal projections of the four T4/T5 subtypes were organized in a single protolayer (**Figure 1H**). By sparsely labeling T4/T5 neurons using MultiColor FlpOut (MCFO) (Nern et al., 2015) with R42F06-Gal4 (Schnell et al., 2012), a driver that by P20 onwards labels all T4/T5 neurons (**Figure S4**), we found that at these early stages individual neurons extend filopodia capable of reaching both Con^−^ and Con^+^ lobula plate domains (**Figure 1I**). Consistent with previous findings, by P40 we could identify two distinct protolayers: one Con^−^, corresponding to neurons specialized in horizontal motion sensing, and another Con^+^ corresponding to vertical motion sensing (Hormann et al., 2020; Kurmangaliyev et al., 2019) (**Figure 1J**). At this stage, the elongating axonal trunk adopts a distinctive “C” shape, from where small filopodia protrude (**Figure 1K**). By P60, all T4/T5 axonal terminals were segregated in four layers (**Figure 1L**). Thus, T4/T5 neurons single layer innervation emerges through the progressive restriction of axonal projections over a 40-hour period. We then assessed the innervation of different T4/T5 postsynaptic partners. We focused on the layer 3 T4/T5c postsynaptic partners LPi3-4, LLPC2, LPC2, Y3-like and the layer 4 T4/T5d partners LPi4-3 and VS neurons, using cell type specific developmental drivers (Chen et al., 2023; Fujiwara et al., 2017; Klapoetke et al., 2017; Mauss et al., 2015). In all cases, we found that at P20 all cell types already innervated the lobula plate (**Figure S5**). We could not assess VS neurons since the HS/VSGal4 driver only labeled these neurons after P60 (**Figure S5**). These results reveal that neurons postsynaptic to T4/T5 innervate the lobula plate before layer segregation, and that layer segregation occurs at a time when both pre- and postsynaptic neurons already innervate the neuropile. We observed that already at P20, all analyzed neurons that in the adult arborize to one or few layers, namely LPC2 and LLPC2, which only arborize in layer 3, and LPi3-4 and LPi4-3, which span layers 3 and 4, already restrict their innervation to the posterior half of the LoP, corresponding to future layer 3 and 4. At this stage, T4/T5 axonal arborizations were not yet restricted to single layers. This suggests that the restriction of T4/T5-postsynaptic partners to specific layers precedes, and is independent of the restriction of T4/T5 axons to specific layers. Lastly, we asked how the development of this layered organization relates to the development of synaptic connectivity. For this, we looked at the expression of early active zone seeding factor Synapse defective 1 (Syd1). Syd1 plays a crucial role in promoting active zone assembly by recruiting presynaptic components (Dai et al., 2006; Owald et al., 2010; Patel et al., 2006), such as the ELKS/CAST family protein Bruchpilot (Brp) (Dai et al., 2006; Fouquet et al., 2009; Wagh et al., 2006), a ubiquitous presynaptic active zone protein. To investigate the endogenous localization of Syd1 in T4/T5 terminals, we crossed a conditional syd1 endogenously tagged GFP reporter (flipase-dependent syd1::frt-stop-frt-GFP) with the T4/T5 driver R42F06-Gal4>uas-flp, while concurrently immunostaining with an anti-Brp antibody. At P20, Syd1 was localized to T4/T5 terminals, but lacked layer organization, while Brp was barely detectable (**Figure 1M**). By P60, Syd1 was organized in four layers, followed by Brp shortly after at P65 (**Figure 1N, O**). This temporal sequence highlights a developmental process in the establishment of synaptic connectivity. T4/T5 neurons and their postsynaptic partners initially undergo a phased progression wherein they organize their processes into specific layers, followed by subsequent phases of synaptogenesis and synaptic maturation. Moreover, the observation that T4/T5-postsynaptic neurons restrict their processes to specific layers before T4/T5 neurons suggests that layer restriction of T4/T5-postsynaptic partners is independent of the process that establishes T4/T5 single-layer innervation.

### T4/T5 subtypes and their postsynaptic partners express matching complements of Beat/Side IgSF CAMs

The expression of matching sets of CAMs in axons and dendrites of synaptic partners may contribute to layer formation (Baier, 2013; Huberman et al., 2010; Millard and Pecot, 2018). To identify CAMs potentially involved in lobula plate development, we used a comprehensive single-cell transcriptional atlas of the developing visual system (Ozel et al., 2021). This dataset covers five developmental time points and the adult, with over 53 clusters matched to known cell types in the connectome, including all T4/T5 subtypes and several of their postsynaptic partners. We focused on the development of the vertical motion sensing network, for which we had available reagents to manipulate gene expression in specific neuronal populations. Virtually all top differentially expressed genes (DEGs) between T4c and T4d are the same as between T5c and T5d (**FIGURE 2A, B**). Although each T4/T5-c/d subtype, and all postsynaptic partners for which the developmental transcriptome is known express many CAMs (**FIGURE 2C**, **S1**), we found that only two pairs (Side-IV/Beat-IIa/b and Side-II/Beat-VI) members of the Beat and Side heterophilic protein interaction network (Li et al., 2017; Ozkan et al., 2013) of interacting CAMs correlated with their common layer innervation (**Figure 2D**, **S1**). Side-IV was uniquely expressed in T4/T5c while Side-II was expressed in T4/T5d, and weakly in T4/T5a of the horizontal system (**Figure 2E**, **S1**, **S6**). For all T4/T5c/d-postsynaptic neurons for which their transcriptome is available (LPi3-4, LPi4-3, TmY14, Y3-like, LLPC1, TmY5a, TmY4, LPC2, LLPC2, LLPC3) (Ozel et al., 2021; Yoo et al., 2023), we found that layer 3 innervating neurons expressed Side-IV’s binding partners Beat-IIa and/or Beat-IIb, while layer 4 neurons expressed Side-II’s binding partner Beat-VI (**Figure 2E**, **S1**, **S6**). Additionally, using a LPTC HS/VS-specific LexA driver in combination with a Beat-VI::T2a-Gal4 gene trap, we found that VS neurons, which are postsynaptic to T4/T5d, also expressed Beat-VI (**Figure S7**). Collectively, these results reveal a remarkable consistency in the expression of binding partners between T4/T5 neurons and their respective layer-specific postsynaptic neurons.

**Figure 2.**
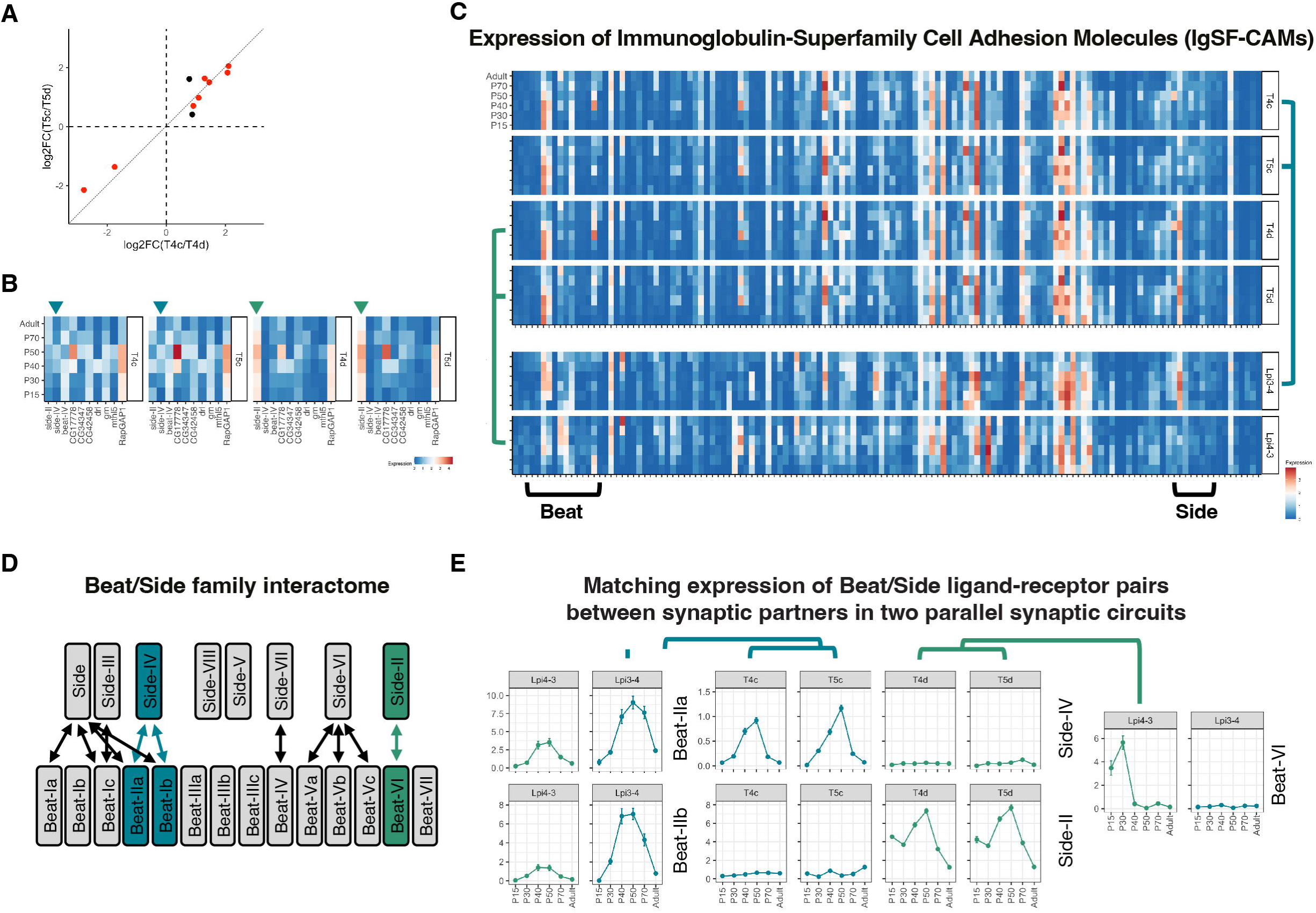
Layer specific expression of matching expression of Beat and Side proteins between synaptic partners. (**A**) Log-fold change for Differentially expressed genes (DEGs) between T4c and T4d (x axis), and T5c and T5d (y axis). See Methods for details and thresholds. (**B**) Average expression level of DEGs (from (**A**)). (**C**) IgSF CAMs in T4/T5 and LPi neurons. Average expression levels are shown for five time points after pupal formation and in the Adult. (**D**) Interactome of Beat/Side families after (Li et al., 2017) (**E**) Line plots with average expression levels of Side-IV, Side-II in T4/T5s and Beat-IIa/b and Beat-VI in LPi neurons.

### Beat and Side display layered localization before T4/T5 axonal segregation and synaptogenesis

Considering these transcriptional expression profiles, we used endogenously-tagged GFP proteins and immunohistochemistry to investigate the localization of Side-IV and its binding partners Beat-IIa and Beat-IIb, as well as Side-II and Beat-VI in the Lobula Plate (**Figure 3**). At P15, when T4T5 neurogenesis ends, Side-II localized in the whole lobula plate while Side-IV was absent (**Figure 3A**). By P25 onwards, both proteins were present in the lobula plate with a characteristic localization pattern. Side-IV was localized only in layer 3 and Side-II in layer 1 and 4, consistent with their T4/T5 subtype specific RNA expression, and effectively dividing the lobula plate in 4 layers (**Figure 3B, C**). At this stage, Beat-IIa showed pan-lobula plate localization, enriched in layer 3 and reduced in layer 4 (**Figure 3D**). Later at P40, both Beat-IIa and Beat-IIb exhibited layer 3-specific localization (**Figure 3E, F, G**), preceding the segregation of T4/T5 axonal terminals in four layers by ~P60 (**Figure 1L**). Unfortunately, our attempts to characterize Beat-VI localization were unsuccessful as the antibodies and Beat-VI protein traps that we generated did not work (data not shown). The precocious layered expression of these molecules, preceding Syd1 and Brp layered expression (**Figure 1N,O**), highlights that layer formation is independent, and takes place before synaptic consolidation. It further suggests that Beat/Side proteins are involved in layer assembly prior to synaptogenesis.

**Figure 3.**
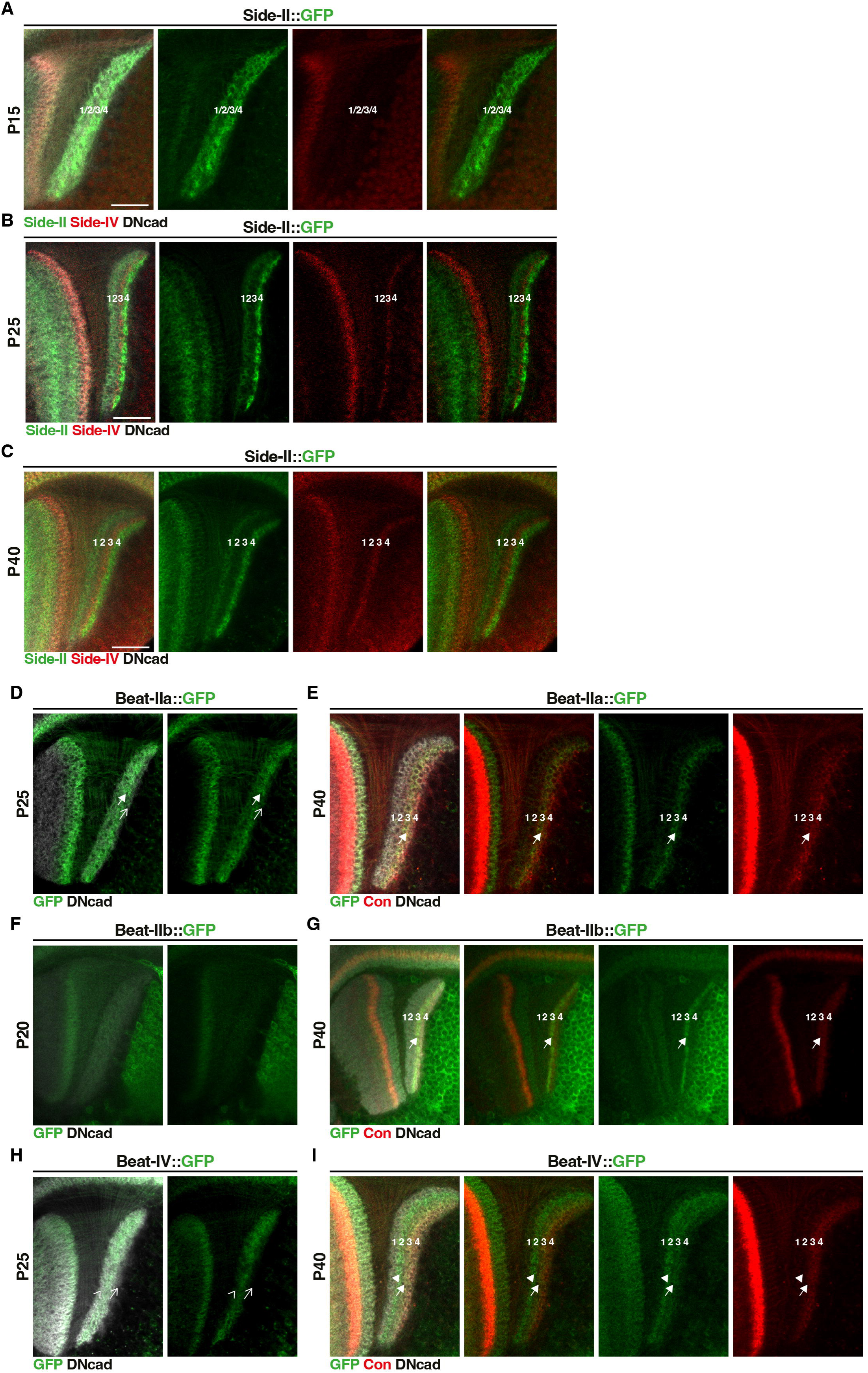
Beat/Side localization in the developing lobula plate. (**A**-**C**) Localization of endogenously tagged Side-II::GFP (green) and Side-IV (Red) proteins during development. (**D-I**) Localization of endogenously GFP tagged Beat-IIa (**D**,**E**), Beat-IIb (**F**,**G**) and Beat-IV (**H**,**I**) during development. Open arrowhead indicates layer 1, closed arrowhead layer 2, closed arrowhead layer 3 and open arrow layer 4). DNcad (grey) labels the neuropiles. Scale bars are 20μm.

### Beats and Sides are required for T4/T5 axons to differentiate between layers c and d

In wildtype, T4/T5 axon terminals form four layers in the lobula plate. In both homozygous *side-II^null^* and *beat-VI^null^*, T4/T5c and d axons were merged, while T4/T5a and b were unaffected (**Figure 4A-C**). This suggests that Side-II and Beat-VI are involved in the same developmental process. Removing Side-II specifically from T4/T5s using R23G12-Gal4 together with side-II RNAi or sgRNA phenocopied homozygous *side-II^null^*, also resulting in T4/T5c/d layer merging (**Figure 4D, E**, **S8**). In contrast, the removal of Beat-VI from T4/T5s did not cause layer 3 and 4 to merge, although a pan-neuronal beat-VI RNAi downregulation did (**Figure 4F, G**). Therefore, Side-II is necessary in T4d/T5d axons, and Beat-VI is necessary in other neurons (including their postsynaptic partners, see below) for the segregation of T4/T5 axonal processes into the same lobula plate layer. Importantly, in T4/T5>side-II RNAi the localization of Side-IV remained restricted to one half of the merged 3/4 layer (**Figure 4A, D**, **S8**). This shows that layers 3 and 4 are merged, but not mixed, and that some sort of layer organization is preserved. I.e., layers 3 and 4 have come together but are not blended, as their individual characteristics are still distinguishable.

**Figure 4.**
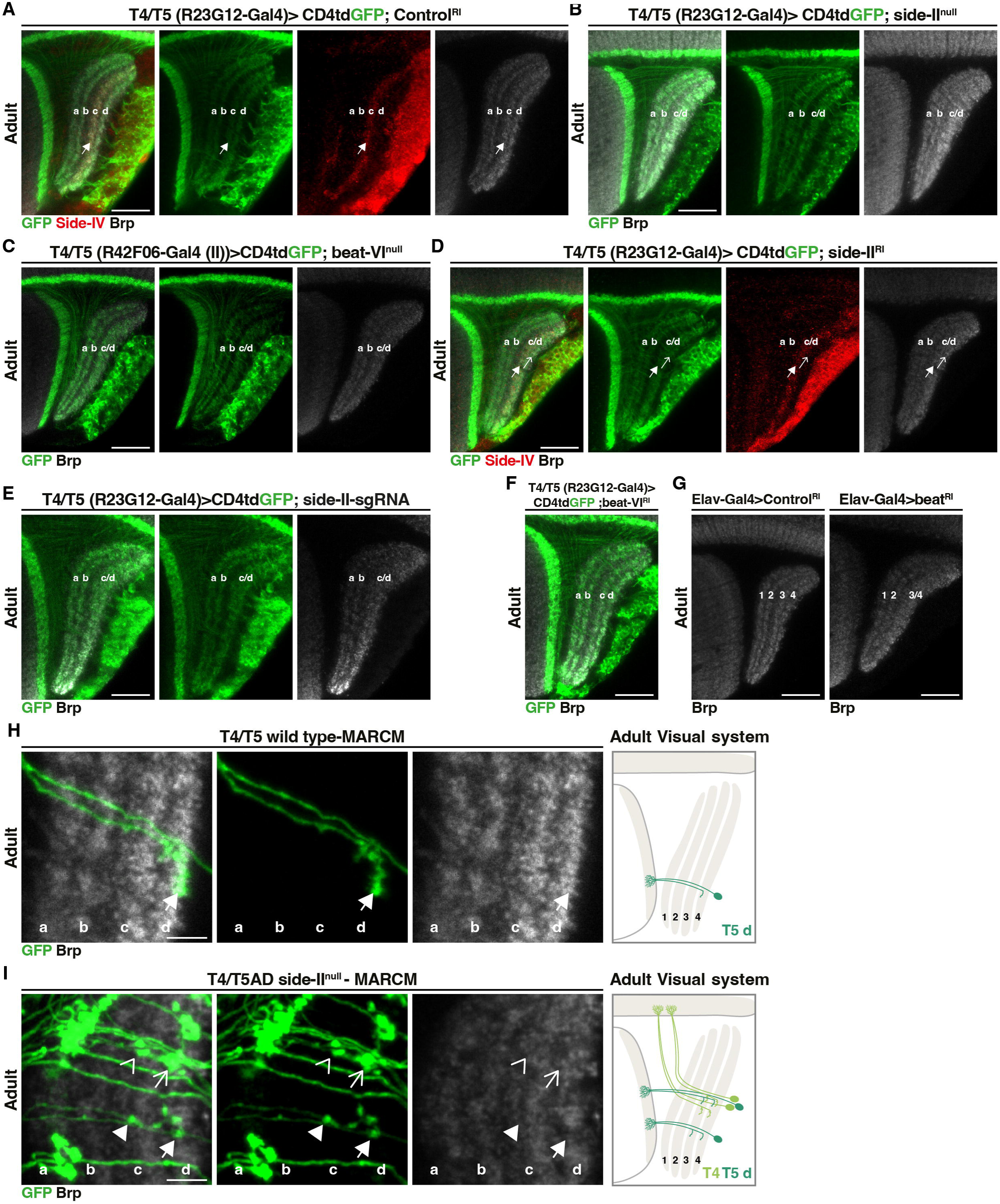
Side/Beat receptor-ligand pair interactions between synaptic partners are required for T4/T5 axons to differentiate between layers c and d. (**A-D**) Morphology of T4/T5s in controls, mutants and upon T4/T5>RNAi specific downregulation. T4/T5 neurons express membrane tethered CD4-tdGFP (green). Side-IV protein (red) (arrow in A and D), Brp (gray) labels the neuropiles. (**A**) Control. (**B**) side-II^null^ homozygous mutants. (**C**) *beat-VI^null^* mutants. (**D**, **E**) Side-II downregulation in T4/T5 neurons with side-II RNAi (**D**) and sgRNA against side-II (**E**). In (**B**-**E**) axons of T4c/T5c and T4d/T5d form a single merged c/d layer. Upon side-II downregulation in T4/T5, Side-IV localization (arrow) remained restricted to half of the merged 3/4 layer (compare A with D). In (D) open arrow points to the half of the merged 3/4 layer where Side-IV is not localized. (**F**) beat-VI^RI^ in T4/T5 neurons. (**G**) Pan-neuronal beat-VI downregulation with Elav-Gal>beat-VI^RI^. (**H-I**) MARCM clones of T4/T5d neurons. (**H**) Wild type T4/T5d subtype neuron innervates layer 4 (closed arrow) (N=6). (**I**) *side-II*^null^ (N=12) d subtype MARCM clones always innervate both layers 3 (arrowheads) and 4 (arrows). Closed arrow/arrowhead identify projections of a single d subtype neuron innervating both layer 3 (closed arrowhead) and layer 4 (closed arrow). All controls for figures 4 and 5 are compiled in Figure S8. Scale bars are 20μm, and 5μm in (H-I).

To better understand how Side-II/Beat-VI interactions contribute for T4/T5d axonal layer innervation, we visualized single T4/T5d *side-II^null^* neurons with Mosaic Analysis with Repressible Cellular Marker (MARCM) (Lee and Luo, 2001) in combination with the T4/T5 R42F06-gal4 that labels all T4/T5 neurons, or T4/T5-a/d-Gal, which only labels subtypes a and d (**Figure S4**). While in the wild type, T4/T5d axons were restricted to layer 4 (**Figure 4H**) (N=6), in *side-II*^null^ MARCM clones, T4/T5d axons always innervated layers 3 and 4 (**Figure 4I**) (N=12). Thus, Side-II is not needed for T4/T5d axons to innervate layer 4, but its activity is required to prevent the innervation of the adjacent layer 3. In summary, side-II functions in T4/T5d neurons, while Beat-VI is essential in T4/T5d postsynaptic partners to restrict T4/T5d synaptic terminals to a single layer. The layer specific localization of Side-IV suggested a similar role in T4/T5c. However, neither side-IV downregulation in T4/T5 using R23G12-Gal together with Side-IV RNAi or sgRNA, nor homozygous side-IV^null^ displayed any obvious T4/T5 layer segregation defects (**Figure 5A-D**, **S8**). T4/T5b and c also express Beat-IV, which is absent from T4/T5d (**Figure S1**, **S6**). Using an available endogenously GFP-tagged Beat-IV reporter, we found that, much like Side-IV, Beat-IV shows layer specific expression from P25 onwards (**Figure 3H, I**). While homozygous beat-IV ^null^ or side-VII^null^ (Beat-IV binding partner) did not show any layer merging phenotype, side-IV^null^, side-VII^null^ double mutants displayed merged T4/T5c/d axonal projections (**Figure 5E,G**). A recent manuscript that also identified a role for Side-II/Beat-VI interactions in lobula plate layer organization is in line with our findings (Yoo et al., 2023). Collectively, we demonstrate that the coordinated expression of receptor-ligand pairs of Beat and Side proteins between synaptic partners establishes parallel synaptic layer organization by restricting T4/T5 axons to specific layers.

**Figure 5.**
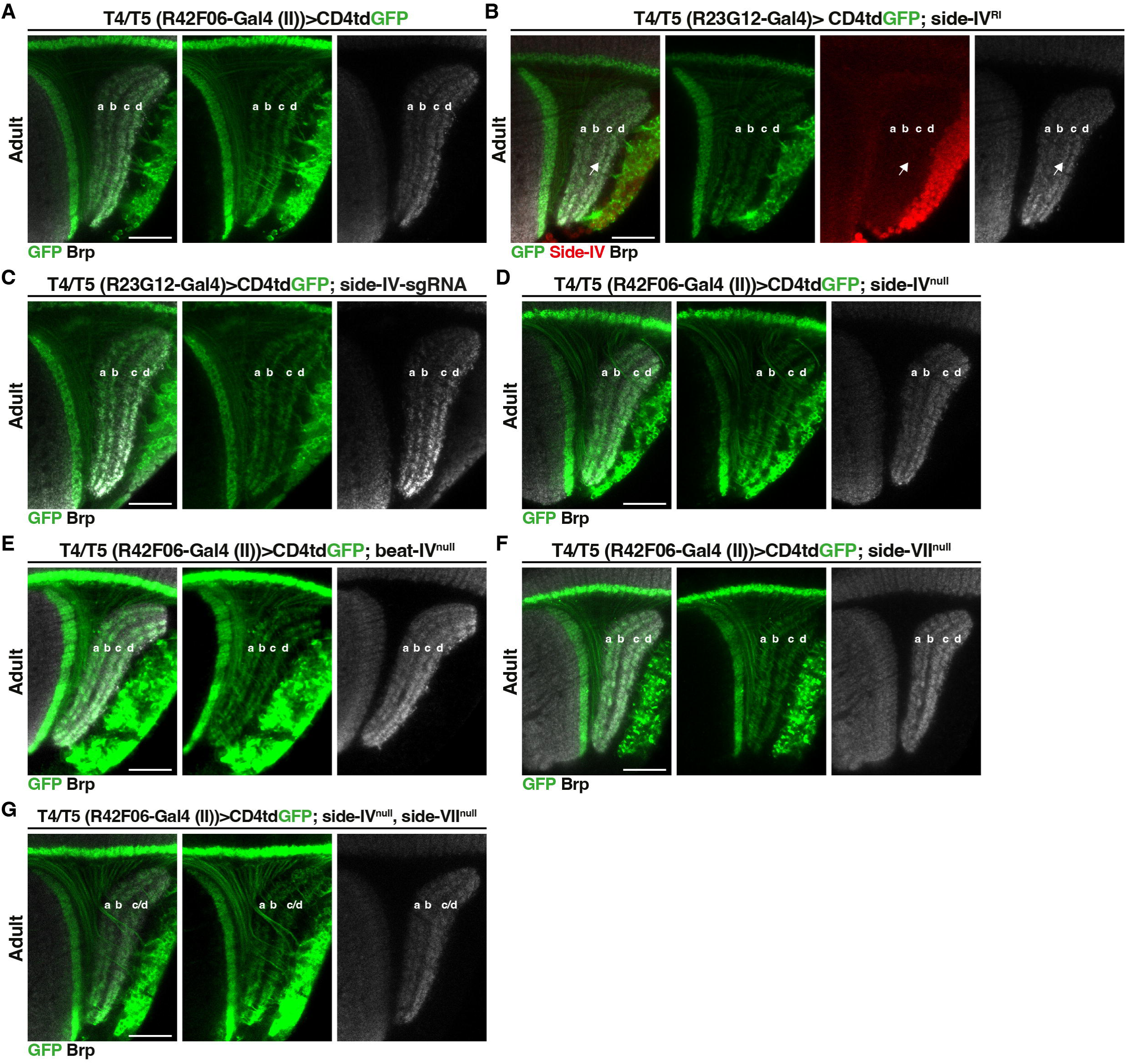
Beats and Sides are required for T4/T5 axons to differentiate between layers c and d. Morphology of T4/T5s in controls, mutants and upon T4/T5>RNAi specific downregulation. T4/T5 neurons express membrane tethered CD4-tdGFP (green). Side-IV protein (red), Brp (gray) labels the neuropiles. (**A**) Wild type control. (**B-C**) T4/T5-specific side-IV downregulation with RNAi (**B**), and sgRNA (**C**). Side-IV is absent from the lobula plate upon T4/T5>side-IV^RI^ (arrow, compare with Figure 4 and Figure S8) (**D-G**) Morphology of T4/T5s in Side-IV^null^ (**D**), Beat-IV^null^ (**E**), Side-VII^null^ (**F**) and Side-IV^null^, Side-VII^null^ double mutants (**G**). T4/T5 layer segregation was impaired in the double mutant but not in the single mutants. All controls for figures 4 and 5 are compiled in Fig S8. Scale bars are 20μm

### Beat/Side interactions are not required to restrict T4/T5-Postsynaptic neurons layered innervation

Next, we sought to assess the role of Beat/Side interactions in the wiring of T4/T5-postsynaptic partners. We focused on the postsynaptic neurons that, together with T4/T5, establish the lobula plate motion opponency circuit: the LPi3-4, LPi4-3 and the LPTC-VS (**Figure 1C-G**, **S2**)). In the wild type, VS dendrites are restricted to layer d, where they receive input from T4/T5d (**Figure 1G**). Likewise, LPi4-3 neurons extend their dendrites into layer d, where they are postsynaptic to T4/T5d neurons. Simultaneously, LPi4-3 neurons project their axons to layer c (**Figure 1F**). A similar circuit exists in layer c, where LPi3-4 neurons extend their dendrites into layer c (**Figure 1E**), acting as postsynaptic partners to T4/T5c neurons (**Figure S2**). These neurons also extend their axons to layer d. First, we downregulated side-II in T4/T5 neurons with R23G12-Gal>side-II RNAi, and concurrently visualized LPi4-3 or VS neurons with cell type specific drivers LexA drivers. Neither LPi4-3 or VS neurons displayed layer innervation defects, and were indistinguishable from the WT (**Figure 6A, B**, **S9**). To further confirm these results, we removed Beat-VI in LPi4-3 neurons using the LPi4-3 developmental driver R38G02-Gal4 (**Figure 6C, D**), while co-expressing the presynaptic marker synaptotagmin::GFP (syt::GFP) (Zhang et al., 2002) to label its axonal processes in layer 3, and the dendritic marker Denmark::RFP (Nicolaï et al., 2010), which strongly localized in dendritic processes in layer 4, and overfilled the neuron. Beat-VI downregulation had no impact on LPi4-3 layer innervation, which remained unaffected and indistinguishable from the wild type (**Figure 6C, D**). Likewise, the downregulation of side-IV in T4/T5 neurons using R23G12-Gal4 did not have any discernible effect on LPi3-4 layer innervation (**Figure S1**). Altogether, we find no evidence for a role for Beat/Side interactions in restricting T4/T5-Postsynaptic neurons layer innervation.

**Figure 6.**
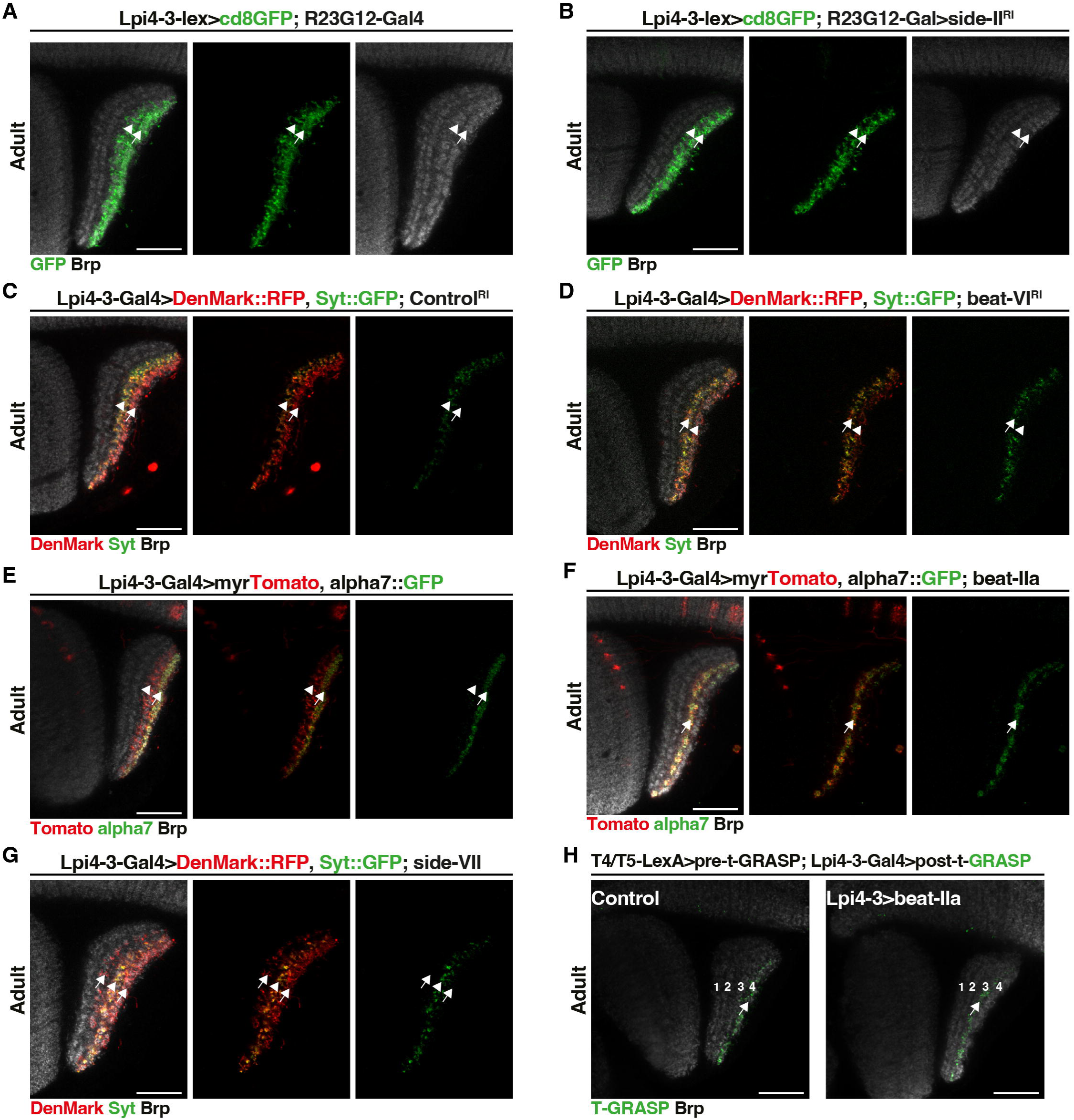
Beat/Side overexpression is sufficient to change layer specific innervation. LPi4-3 visualized with R38G02-LexA or -Gal4 in the wild type and upon manipulation of Beat/Side interactions. In all panels, arrow points to dendritic terminals and arrowhead to axonal terminals. Brp (grey) labels the neuropiles. (**A-B**) LPi4-3 neurons expressing membrane tethered CD8tdGFP (green) in the wild type (**A**), and upon side-II^RI^ mediated downregulation in T4/T5 neurons (**B**). (**C-D**) Wild type LPi4-3 neurons expressing Syt::GFP in the axonal terminals (arrowhead) in layer 3, and Denmark::RFP that labelled the neurons, with stronger expression in the dendrites (arrow) in layer 4 (**C**), and upon beat-VI^RI^ downregulation (**D**). (**E-F**) Wild type LPi4-3 expressing the nicotinic acetylcholine receptor subunit Dα7, which localized to LPi4-3-cholinergic synapses (Ammer et al., 2023) (**E**), and upon beat-IIa overexpression (**F**). Overexpression of Beat-IIa in LPi4-3 induced the innervation of only layer 3, where the neurons developed Dα7^+^ dendritic processes (arrow). (**G**) Overexpression of Side-VII in LPi4-3 neurons expressing Syt::GFP and Denmark::RFP. Side-VII overexpression promoted the development of dendritic processes in layer 2 (arrow), where the partner of Side-VI, Beat-IV, is strongly expressed in T4/T5 axons. LPi4-3 neurons maintained their dendritic and axonal innervations in layers 4 and 3, respectively. (**H**) t-GRASP activity-independent GRASP between T4/T5 neurons (R42F06-Lexa) and LPi4-3 (R38G02-Gal4) neurons in the wild type (left), and upon beat-IIa overexpression in LPi4-3 neurons (right). Scale bars are 20μm.

### Beat/Side overexpression is sufficient to change layer specific innervation

To test if Beat/Side ectopic expression is sufficient to promote changes in layer innervation, we first used our LPi3-4 developmental Split-Gal4 driver (**Figure S5**) to overexpress beat-VI. This was sufficient to re-direct practically all the neurons dendritic processes to layer 4 (**Figure S10**), where the partner of Beat-VI, Side-II, is strongly expressed in T4/T5 axons. We then overexpressed beat-IIa in LPi4-3, while co-expressing the nicotinic acetylcholine receptor subunit Dα7, which localizes to LPi4-3-cholinergic synapses with T4d/T5d neurons (Ammer et al., 2023) (**Figure 5E**). Beat-IIa overexpression induced LPi4-3 innervation of only layer 3, where the neurons developed Dα7^+^ dendritic processes (**Figure 5F**). In contrast, when we overexpressed side-VII, Lpi4-3 extended its dendritic processes to layer 2, where the partner of Side-VII, Beat-IV, is strongly expressed in T4/T5 axons, while maintaining their dendritic and axonal innervations in layers 4 and 3, respectively (**Figure 5G**). These results demonstrate that the overexpression of beat/side genes alone is sufficient to modify layer innervation patterns. Moreover, the distinct layer innervation phenotypes between LPi4-3>side-VII, characterized by the retention of dendrites in layer 4, and LPi4-3>beat-IIa, where such dendritic preservation is absent, highlight variations in adhesive interactions between neurons. This difference suggests that the preferential adhesion among neurons may hinge on factors such as the intensity of specific beat-side interactions, other heterophilic IgSF cell adhesion molecules (Xu et al., 2022), the complete array of CAMs expressed by the neurons, and the distance between the axons and the target layers.

### Beat/Side dependent changes in layer innervation are sufficient to change connectivity

Having established that Beat-IIa overexpression in layer d-postsynaptic neurons shifts their innervation to layer 3, with concomitant neurotransmitter receptor expression in layer 3-innervating dendrites, we proceeded to examine whether this altered receptor expression resulted from ectopic synapses with T4/T5c neurons. To test this hypothesis, we used t-GRASP to probe for synaptic connectivity between Lpi4-3 and T4/T5 neurons. T-GRASP is an activity-independent, targeted GRASP, where one of the split-GFP fragments is targeted to presynaptic terminals and the other to dendrites, such that GFP reconstitution only occurs at synaptic sites (Shearin et al., 2018). In the wild type, robust t-GRASP signals were evident in layer 4, corresponding to the presynaptic localization of T4/T5d neurons with LPi4-3 (**Figure 5H**) (N=4). However, upon the overexpression of beat-IIa in LPi4-3, we observed t-grasp signals in layer 3 and not in 4 (**Figure 5H**) (N=4). This indicates that Beat/Side-dependent alterations in layer innervation coincide with changes in synaptic specificity between parallel circuits.

### Beat and Side promote adhesiveness prior to synapse formation

We have shown that Beat/Side ectopic expression is sufficient to promote changes in layer innervation, with the development of ectopic synapses with non-canonical synaptic partners. The next question to ask is if Beat/Side mediated adhesiveness during development occurs prior to or concurrently with synapse formation. To answer this question, we looked at the layer innervation of LPi4-3>beat-IIa during development. As early as P25, the projections of LPi4-3>beat-IIa in the lobula plate are confined to the prospective layer 3. This contrasts with the wild type, where innervation spans half of the lobula plate neuropile, encompassing layers 3 and 4 (**Figure 7**). Notably, this occurs ~40 hours before the onset of Syd1 and Brp layered expression. These observations suggest that Beat/Side interactions may contribute to adhesiveness between synaptic partners prior to synapse formation.

**Figure 7.**
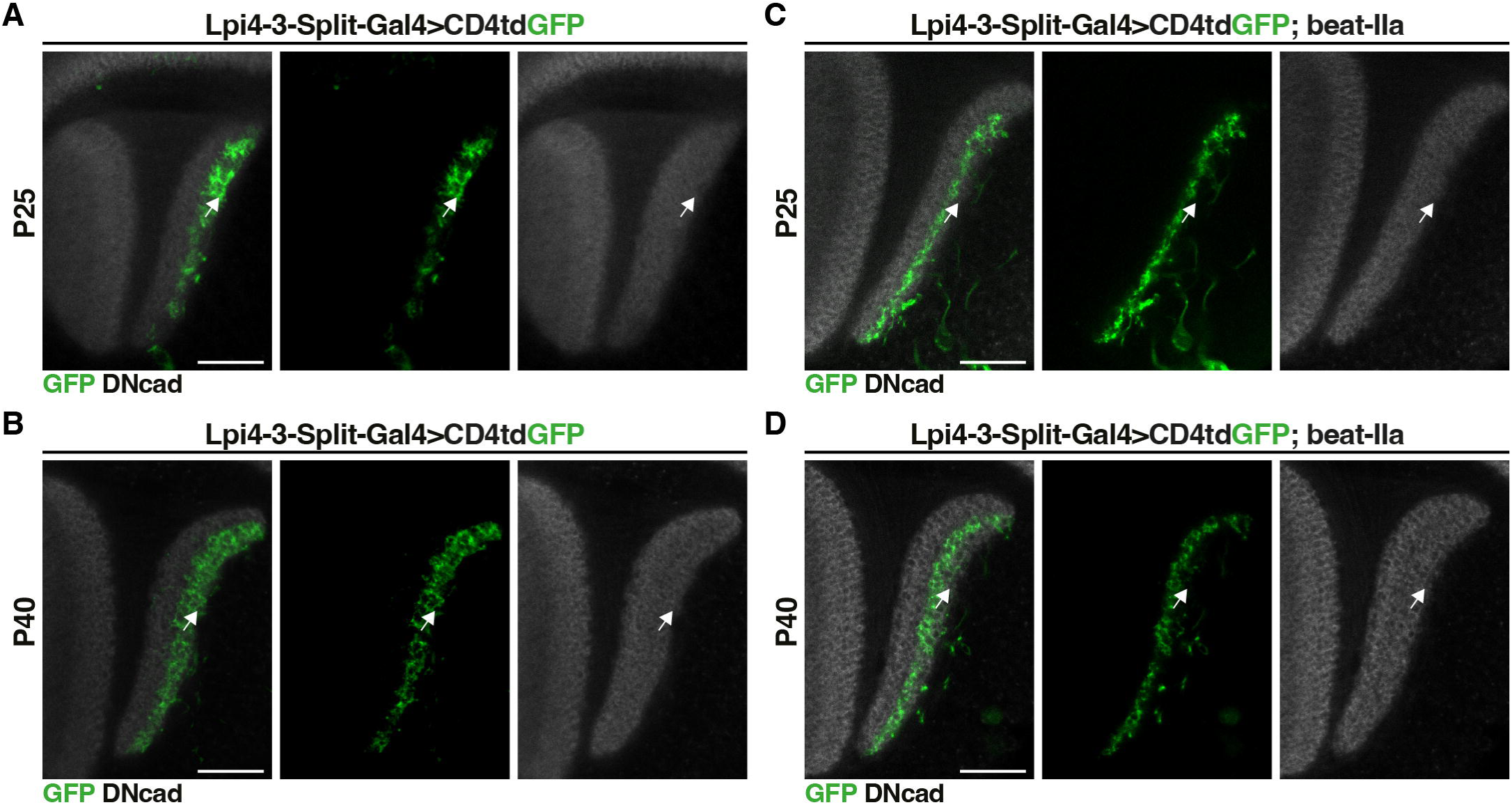
Beat/Side interactions mediate cell adhesion before synaptogenesis. LPi4-3-Split-Gal4 driving membrane tethered CD4-tdGFP (green). DNcad (grey) labels de neuropiles. LPi4-3 lobula plate innervation in the wildtype. Arrow point to the right edge of the lobula plate where layer 4 develops. (**A-B**) wild type. (**C-D**) LPi4-3 overexpressing beat-IIa. Beat-IIa overexpression promotes LPi4-3 exclusive layer 3 innervation as early as P25, ~40h before synaptogenesis. Scale bars are 20μm.

## DISCUSSION

The organization of neuronal processes into layers emerges through the action of long-range axon guidance and topographic mapping mechanisms, which bring axons and dendrites in spatial proximity, restricting the number of potential synaptic partners. Once neurons are in close proximity, extracellular short-range guidance cues and cell surface adhesion may contribute to synaptic specificity. Thus, synaptic specificity is the end result of a multistep developmental process that sequentially restricts the pool of possible partners. However, the molecular and cellular mechanisms governing the organization of neuronal processes in distinct layers, and how these contribute to the development of robust synaptic connectivity remain unclear. We tackled this problem by studying the development of the lobula plate neuropile, whose layer-specific arrangement of neuronal connections sustains the parallel processing of different visual motion cues. We found that each subtype of T4/T5 axons, along with their corresponding layer-specific synaptic partners, express matching Beat/Side receptor-ligand pairs. Our findings reveal that these molecules are not needed for T4/T5 neurons to innervate their default layer. Instead, our results suggest that these receptor-ligand pairs promote T4/T5 axons single layer innervation by biasing cellular adjacency to postsynaptic partners before synaptogenesis. We further propose that, in each layer, synaptic specificity depends on competitive interactions involving shared Beat/Side expression among all postsynaptic partners, each competing to establish connections with T4/T5 neurons.

### Beat/Sides promote T4/T5 axons single layer innervation by biasing cellular adjacency to postsynaptic partners

We have demonstrated the essential role of T4/T5 neurons in the development of the lobula plate, as the neuropile fails to form in their absence. Simultaneously, our findings reveal that the organization of T4/T5 axons into distinct layers relies on Beat/Side-mediated interactions between T4/T5 axons and their postsynaptic neurons. Taking layer 4 as an example, removing Beat-VI, or ectopically expressing Beat/Sides in specific post-synaptic partners (Lpi4-3 neurons), did not affect layer formation. Contrarily, in side-II^null^ or homozygous beat-VI^null^, layers 3 and 4 were merged, because T4/T5d axons now innervate both layers 3 and 4. These observations reveal that the segregations of layers 3 and 4 depends on the widespread expression of Beat-VI in many, if not all, T4/T5d-postsynaptic neurons to sustain the innervation of T4/T5d to a single layer (layer 4). The role of T4/T5 neurons in the development of the lobula plate can be conceptualized as a reciprocal relationship. On one hand, T4/T5 neurons act as essential anchors for the formation of the neuropile, serving as a structural foundation for other neurons to join in a Beat/Side independent manner. On the other hand, the correct segregation of T4/T5 axons into distinct layers depends on their Beat/Side-mediated spatiotemporal coordinated interactions with postsynaptic neurons.

During the writing of our manuscript, a study was published, proposing that removal of Side-II/Beat-VI interactions induces layer c/d fusion and leads to inappropriate targeting of both T4/T5 axons and the dendrites of their respective postsynaptic partner (Yoo et al., 2023). Our observations support a role for Side-II/Beat-VI interactions mediating T4/T5 axonal layer innervation, for which we offer a mechanistic explanation. However, the suggestion that removal of Side-II/ Beat-VI interactions induces layer fusion, while also mediating postsynaptic partners layered innervation, is difficult to reconcile with our observations. These include the characterization of the layered organization of postsynaptic neurons prior to T4/T5 layered axonal organization, and before the Beat/Side-layered localization. In addition, the presence of localized Side-IV protein expression in Side-II mutants, and the detailed characterization of postsynaptic neuron morphologies using axonal and dendritic markers upon removal of Side-II/ Beat-VI interactions, neither of which support this notion. Both studies, together with a recent pre-print (Osaka et al., 2023), identify Beat/Side-mediated interactions between synaptic partners, and expand the repertoire of IgSF protein families implicated in circuit development in *Drosophila*.

### From cellular adjacency to synaptogenesis

The process of synapse formation is highly resilient (Hassan and Hiesinger, 2015). In either vertebrates and invertebrates, the dysregulation of proteins contributing to synaptic specificity (Krishnaswamy et al., 2015; Shen and Bargmann, 2003; Shen et al., 2004) or synaptic organization (Chen et al., 2017; Mosca et al., 2012; Robbins et al., 2010; Südhof, 2017) does not necessarily prevent synaptogenesis. In addition, when neurons do not have their usual counterparts or when they connect to different brain regions, they establish ectopic synapses with alternative partners (Bekkers and Stevens, 1991; Cash et al., 1992; Duan et al., 2014; Shen and Bargmann, 2003; Zhang et al., 2022). These observations support the notion that synapse specificity and synapse assembly are separable (Kurshan and Shen, 2019). An important result from our study, and that of Yoo et al. (Yoo et al., 2023), is the observation that Beat/Side interactions are not needed for T4/T5 axons to innervate their default layer and establish synapses. In the fly optic lobe, L2 and L4 make reciprocal synapses (Rivera-Alba et al., 2011). In a recent preprint, Osaka et al. show that Side-IV in L2 neurons interacts with Beat-IIa/b in L4 neurons to promote adjacency (Osaka et al., 2023). Additionally, the authors provide compelling evidence that Side-IV can interact in cis with the synaptic scaffolding protein Kirre, a member of the irre cell recognition module (IRM) family, and recruit Syd-1 through the intracellular PDZ domain of either Side-IV or Kirre. Together with our findings, the evolving framework provides a cellular and molecular mechanism through which the same ligand-receptor pair influences inter-cellular adjacency with a synaptic partner and recruits synaptic seeding factors to influence synapse induction: Initially, Beat/Side mediated inter-cellular interactions influence the likelihood of adjacency before synaptogenesis through extracellular, domain-based preferential adhesion mechanisms (our study). Subsequently, Beat/Side can enhance the likelihood of synapse formation by aiding in the recruitment of seeding factors to sites of adjacency (Osaka et al., 2023). Implicit in the concept of biasing the recruitment of synaptic seeding factors is the assumption that these factors are a finite resource. Supporting evidence comes from live imaging studies of the axons terminals of R7 photoreceptors at the time of synaptic partner choice (Ozel et al., 2019). Here, the formation of one synapse at a time suggests that the number of synapses formed is determined by the presynaptic neurons, and limited by the availability of synaptic seeding factors. Furthermore, loss of R7’s main postsynaptic partner Dm8 affects neither serial synapse formation nor the total number of R7 synapses, albeit now occurring with non-canonical postsynaptic partners (Kiral et al., 2021). Also, in L4 neurons, the absence of DIP-β (of the Dip/Dpr interacting families) does not result in a loss of synapses at the typical DIP-β localization site (Xu et al., 2019). Instead, its loss promotes the formation of ectopic synapses in a neighbouring region where DIP-β is not typically found. This observation prompted the authors to first propose that neurons have the capacity to synapse with several cell types and that synapse specificity is achieved by establishing a preference for specific partners.

Beat/Side represents one of the various families of heterophilic-interacting cell adhesion molecules that can contribute to adhesiveness between neurons. These molecules exhibit broad expression throughout the developing visual system system (Ozel et al., 2021; Tan et al., 2015; Yoo et al., 2023) and show layered localization in other optic lobe neuropiles (**Figure 3**), indicating a potential contribution to circuit development on a broader scale. A standing challenge in the field is to understand what the specific contribution of various CAMs is to distinct cellular behaviors, and how each participates in the process of building synaptic specificity. Future 4D studies exploring the timing and subcellular distribution of different CAMs, and investigating their roles in processes such innervation dynamics, adjacency and synaptogenesis, will help addresssing it.

### Competitive adhesive dynamics among synaptic partners sustain the development of robust connectivity

Our study suggests that the establishment of inter-neuronal adjacency is intricately tied to a competitive dynamic fostered by the shared expression of the same Beat partner among postsynaptic partners within a layer, corresponding to the Side expressed the respective T4/T5 presynaptic neurons. We propose a previously overlooked function of layered organization in shaping the adhesion between neurons, and ultimately the synaptic specificity of neural circuits. Within the physical spaces of each layer, shared molecular context establish a competitive milieu contributing to synaptic specificity by influencing the adhesion dynamics of each neuron. This perspective offers valuable insights into the cellular mechanisms responsible for establishing layered organization. Furthermore, it presents a general framework for undestanding how the process of layer assembly actively contributes to the development of robust neural circuits in the brain.

## Supporting information

Resource Table

## Acknowledgements

We would like to thank, Larry Zipursky, Juyoun Yoo, Mark Dombrovski and Neset Ozel for helpful discussions, and Jesus Pujol, Nikos Konstantinides, Hernán Lopez-Schier, Robin Hiesinger and Claude Desplan for comments on the manuscript. We thank the fly community, especially Takashi Suzuki, Axel Borst and Claude Desplan for gifts of plasmids, antibodies and fly stocks. Also, the MCD/CBI fly facility for help in the generation of transgenic lines. This work was supported by a Fondation Bettencourt Schueller grant (“Building neural networks for motion vision”) under the CNRS-INSERM-ATIP-Avenir 2019 program to F. P-T. N. F. was supported by a FRM postdoctoral fellowship (SPF202110013946). V. d. S. X. was supported by a PhD fellowship (2020.06759.BD) from the Portuguese Foundation for Science and Technology. M. T. was supported by a PhD fellowship from the Paul Sabatier University BSB Doctoral School, and by a FRM PhD fellowship (FDT202304016913). This work was supported in part by a grant from the National Institutes of Health (NIH) (NIH NRSA F32EY032750) to Y.-C.D.C.

## RESOURCE AVAILABILITY

### Lead contact

Further information and requests for resources and reagents should be directed to and will be fulfilled by the lead contact, F.P-T.

### Materials availability

Newly generated mutant alleles and split-Gal4 fly lines are available upon request.

## EXPERIMENTAL MODEL DETAILS

### Fly husbandry and genetics

Flies were kept on standard cornmeal medium at 25°C, 12-hour light/dark cycles. All RNAi experiments were run with UAS-Dcr2.D at 29°C. No differences were observed in neuronal morphologies between control flies kept at 25°C and 29°. All transgenic flies used in this study are described in Resource Table S1. Full genotypes of flies in each experiment are described in Table S2. Male and Female pupae and 3–10 day old adult flies were analysed. No differences were observed between sexes.

### Generation of LPi3-4-Split-Gal

LPi3-4-Split-Gal was built as described in (Chen et al., 2023). LPi3-4-Split-Gal is built from Tj-T2a-Gal4vp16 + TkR86C-t2a-Gal4DBD, which co-expression we found to be specific to LPi3-4 and another unidentified neuron that projects in the Medulla and Lamina neuropiles. Based on the co-expression of Tj and TkR86C, LPi3-4 corresponds to cluster 120 in the transcriptional atlas (Ozel et al., 2021).

### Generation of FlpTag lines

Side-II::FlpTag and Beat-IIa::FlpTag constitutive lines where built by injecting the FlpTag plasmid (a gift from A. Borst) in Side-II MIMIC or Beat-IIa MIMIC as previously described (Fendl et al., 2020). Embryo injections were done at the MCD/CBI Fly facility.

### Cas9-mediated mutagenesis of side-II

To generate a side-II^null^, we introduced a frameshift mutation on Side-II allele using the TRiP-KO line (BDSD 82048) in flies carrying FRT40.

## METHODS DETAILS

### Synaptic connectivity

Synaptic connectivity data corresponds to 40 representative T4/T5 neurons (five instances of each T4/T5 subtype), after (Shinomiya et al., 2022).

### Heatmap of synaptic weights

For each representative neuron, we summed the total number of synapses between each T4/T5 neurons and all synaptic partners of the same type. This data was plotted as a heatmap of synaptic weights between T4/T5 subtypes and postsynaptic partners (**Figure S1**). After (Shinomiya et al., 2022).

### Connectome graph

Synaptic weights between each T4/T5 and each postsynaptic partner were averaged to generate a cell type-level connectome graph. Only connections with more than 10 synapses with a single T4/T5 neuron were plotted. After (Shinomiya et al., 2022).

### Transcriptome analysis

Two transcriptional atlas of the *Drosophila* visual system were independently analyzed (Ozel et al., 2021; Yoo et al., 2023). For figure assembly, data shown is from Ozel et al. 2021 except for Figure S1D, F. which is from Yoo et al.. Single-cell analysis was performed using Seurat (V4.1.1) (Butler et al., 2018). All functions were used with default parameters unless otherwise indicated.

### Visualization of gene expression patterns

Expression patterns were visualized using average expression levels for each cell type and time point. Averaging was performed in non-log space for the original normalized expression values. For the line plots, we used expression values in linear scale. The list of all IgSF proteins was obtained from FlyXCDB (Pei et al., 2018) and (Vogel et al., 2003)

### Differential gene expression analysis

We identified differentially expressed genes between T4 and T5 neurons of subtypes c and d using default parameters. Analysis was performed at P40.

### MARCM Clones

MARCM clones were generated and visualized by immunofluorescence staining as previously described (Pinto-Teixeira et al., 2018). Wild type clones were obtained by crossing hsFlp122, Tubgal80 Frt19; Uas-CD8::GFP; T4/T5-Gal4 (R42F06) X Frt19. Heat shock was induced for 20 min at 37°C in L3. side-II^null^ MARCM clones were generated by crossing hsFlp122; Tubgal80 Frt40; T4/T5ad-Gal4 (this study), uas-cd4tdTOMATO X; side-II^null^ FRT40;. Heat shock was induced for 4-10 min at 37°C in L3.

### Multicolor Stochastic Labelling (MCFO)

“Multicolor FlpOut” (MCFO) labelling was carried out as described before (Nern et al., 2015). Briefly, FLP recombinase expression was used to excise FRT-flanked transcriptional terminators from UAS reporter constructs carrying HA, V5 and FLAG epitope tags. Pupae carrying LLPC2/LPC2-Gal4 and MCFO-1 were raised at 25°C and heat shocked at 37°C for 15min. Immunohistochemistry was performed as per below.

### Immunohistochemistry for confocal microscopy

Pupal brains were dissected in ice-cold 1XPBS, and adult brains in ice-cold Schneider’s Drosophila Medium. Brains were then fixed in 4% Formaldehyde at room temperature for 20 min. Brains were then rinsed 3X times in PBX (1% PBS, 0.3% triton) and left to wash for at least 2 hours and then incubated in primary antibody solution overnight at 4°C for one or two days. Brains were then rinsed 3X times and left to wash for at least 2 hours and then incubated in secondary antibody solution in PBX overnight (with added 2% horse serum) at 4°C. Brains were finally rinsed 3X and left to wash overnight in PBX and mounted in Slowfade. For further confocal imaging, brains were mounted vertically, so that the Z-axis in the final volume corresponds to the D-V axis of the brain.

### Antibody information

Primary antibodies and dilutions used in this study:

rat anti-Side-IV (1:100); guinea pig anti-Brp (1:30); mouse anti-nc82 (1:20), mouse anti-Connectin (1:20) rabbit anti-GFP (1:500); chicken anti-GFP (1:500); sheep anti-GFP (1:500); Phalloidin 405(1:200); rabbit anti-RFP (1:40000); chicken anti-V5 (1:500); rat anti-Dncad (1:20)

Secondary antibodies used at 1:200/1:500. Please view the resource table for additional information and providers.

### Confocal microscopy

Images were acquired using a Leica SP8 confocal or Zeiss LSM 780 using a 40X oil objective. When acquired, Z-stacks sections were of 1–3 μm intervals.

### Image processing

Images were processed with Fiji software package (Schindelin et al., 2012). Image processing, when applied, was limited to correct global images brightness, global background subtraction and Despeckle. To allow the full visualization of neuronal projections of adult MARCM Clones in a 2D image, stacks are shown as maximum intensity Z-projections. Images were compiled in Illustrator (CC2024).

### Quantification and Statistical Analysis

Control or experimental specimens of the correct stage and genotype were randomly and independently selected from a pool of animals. Unless otherwise noted, we examined at least 10 brains for every genotype. In the main text “N” represents the number of samples analysed. Data acquisition was not performed blinded as samples were identified based on their genotypes that were not limited in repeatability. Data analysis was performed blinded.

---

**Figure S1.**
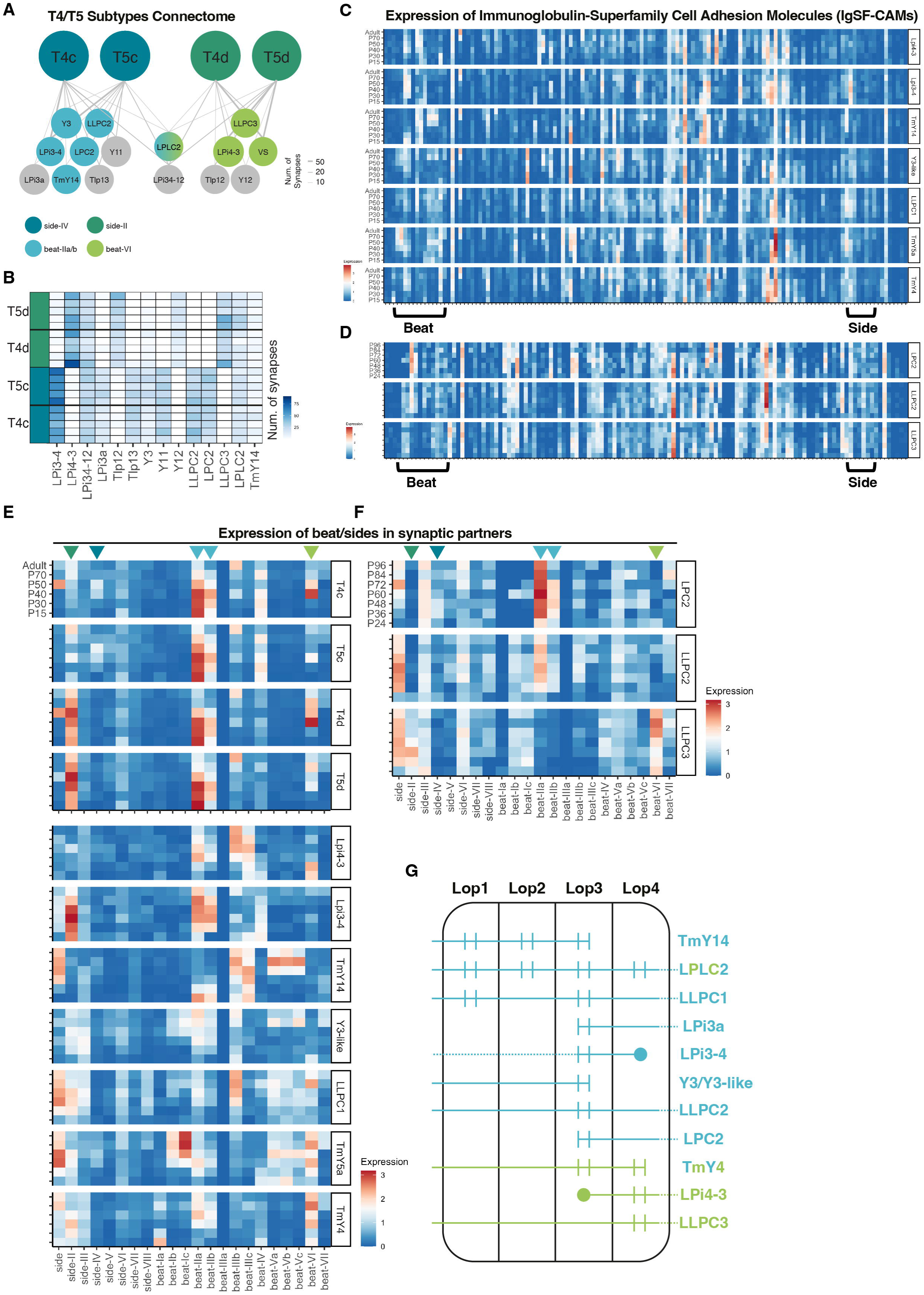
T4/T5 neurons connectome. (**A**) The connectome of T4/T5c and d subtypes with more than 10 synapses with a single T4 or T5 neurons. Color code reflects matching expression of beat and side genes between synaptic partners based on developmental transcriptomes, or Intersectional genetics (for VS neurons). (**B**) Connectomes of five neurons for each T4 and T5 subtype with postsynaptic partners from (**A**). T4 and T5 neurons of the same subtype synapse with the same set of postsynaptic neurons. (**C, G**) All annotated IgSF CAMs (**C, D**) and beat/side (**E, F**) expression in all T4/T5-postsynaptic partners with known developmental and adult transcriptomes. (G) Lobula plate connectivity of the same set of T4/T5c/d-postsynaptic partners. Parallel bars represent synapses with layer specific T4/T5 subtypes. (A), (B) and (G) after (Shinomiya et al., 2022). scRNAseq transcriptome in (C) and (E) from the (Ozel et al., 2021) dataset; and in (**D**) and (**F**) from the (Yoo et al., 2023) dataset.

**Figure S2.**
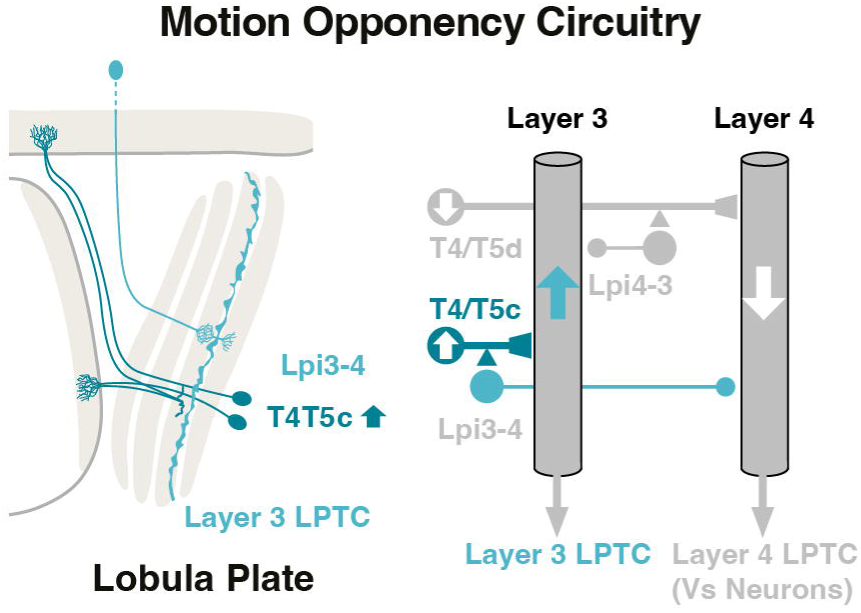
LPi3-4 Motion Opponency Circuit.

**Figure S3.**
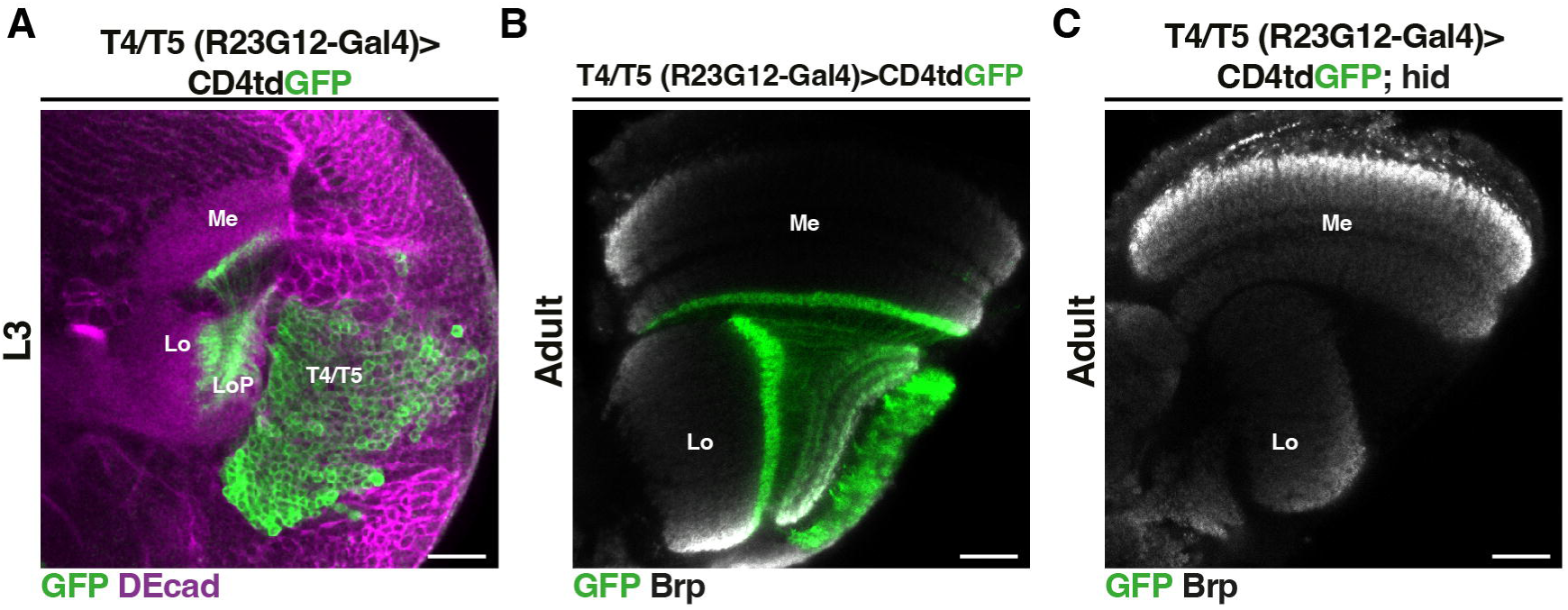
T4/T5 neurons are needed for the development of the lobula plate neuropile. T4/T5 (R23G12-Gal) expression in L3 (**A**) and in the adult (**B**). (**C**) R23G12-Gal4 mediated overexpression of the proapoptotic gene *hid* induces T4/T5 neurons death. The lobula plate neuropile is absent. Abbreviations: Medula (Me), Lobula (Lo), Lobula Plate (LoP). Scale bars are 20μm

**Figure S4.**
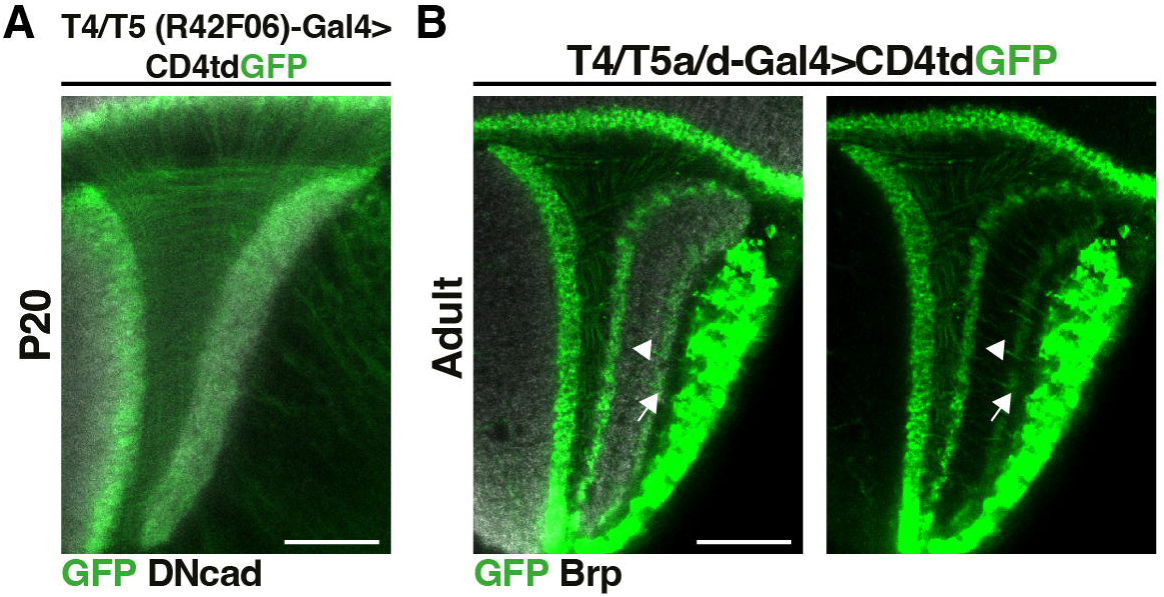
T4/T5-Gal4 drivers characterization. (**A**) T4/T5 (R42F06-Gal4) driving the expression of membrane tethered CD4tdGFP at P20 (green). (**B**) T4/T5-a/d-Gal4 driving the expression of membrane tethered CD4tdGFP T4/T5A/D neurons in the adult (green). Arrow points to layer 4, arrowhead to layer 3. DNCad (grey) in (A) and Brp in (B) label the neuropiles. Scale bars are 20μm

**Figure S5.**
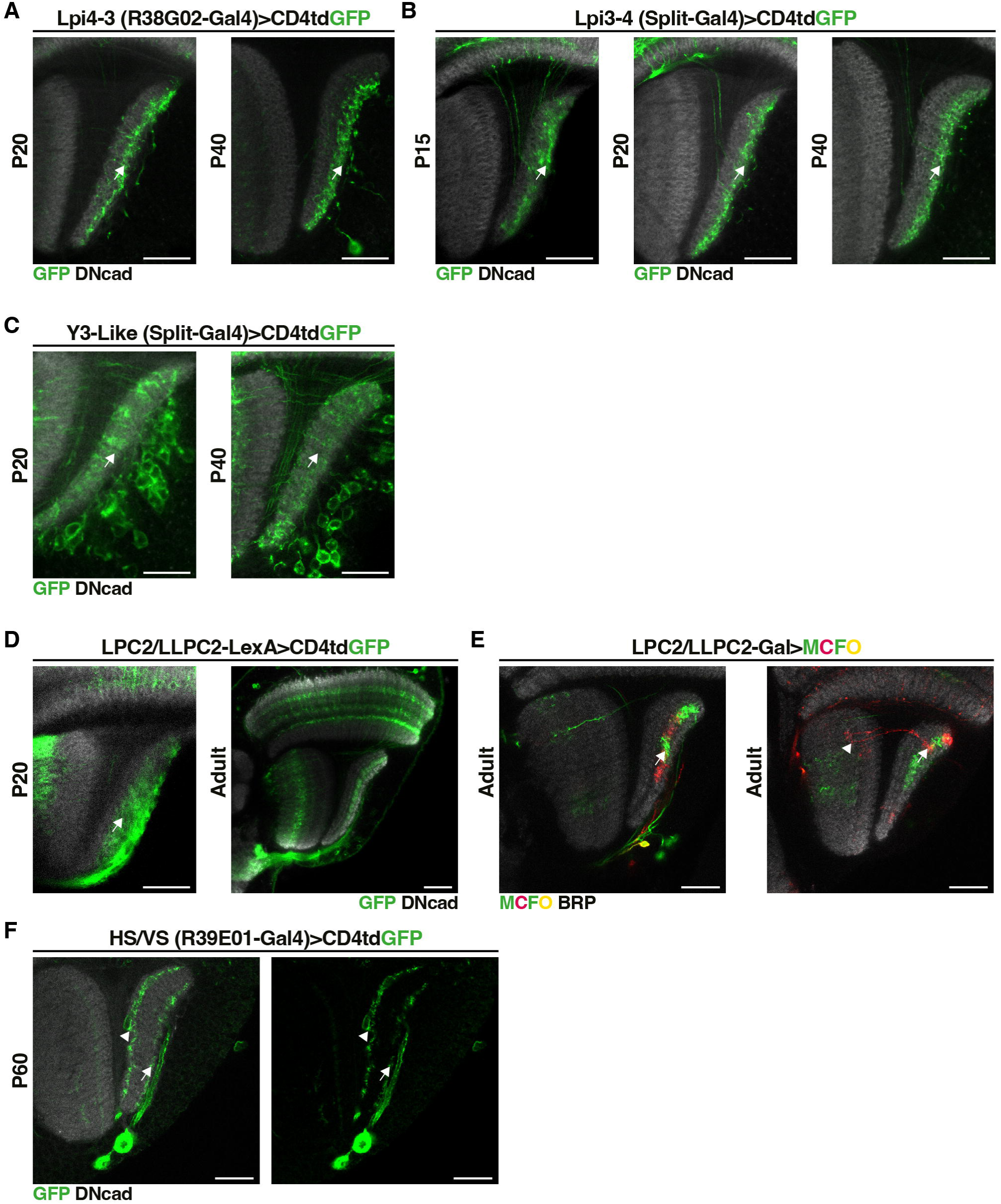
T4/T5-Postsynaptic partners lobula plate innervation during development. Expression of cell type specific Gal4 drivers expressing membrane tethered cd4tdGFP (green). Lobula plate innervation for LPi3-4 (R39G09-Gal4) (**A**), LPi4-3 (Split-Gal4) (**B**), Y3-like neurons (Split-Gal4) (**C**) at P20 and P40, and LPC2/ LLPC2 (**D**) at P20 (left) and adult (right). (**E**) MCFO sparse labeling with LPC2/LLPC2-Gal4 in the Adult. LPC2/LLPC2-Gal4 expression in the lobula plate is labels LPC2 (arrow) and few LLPC2 (arrowhead) neurons. (**F**) HS/VS (R39E01-Gal4) at P60. DNcad and BRP (grey) label the neuropiles during development and in the adult, respectively. Scale bars are 20μm

**Figure S6.**
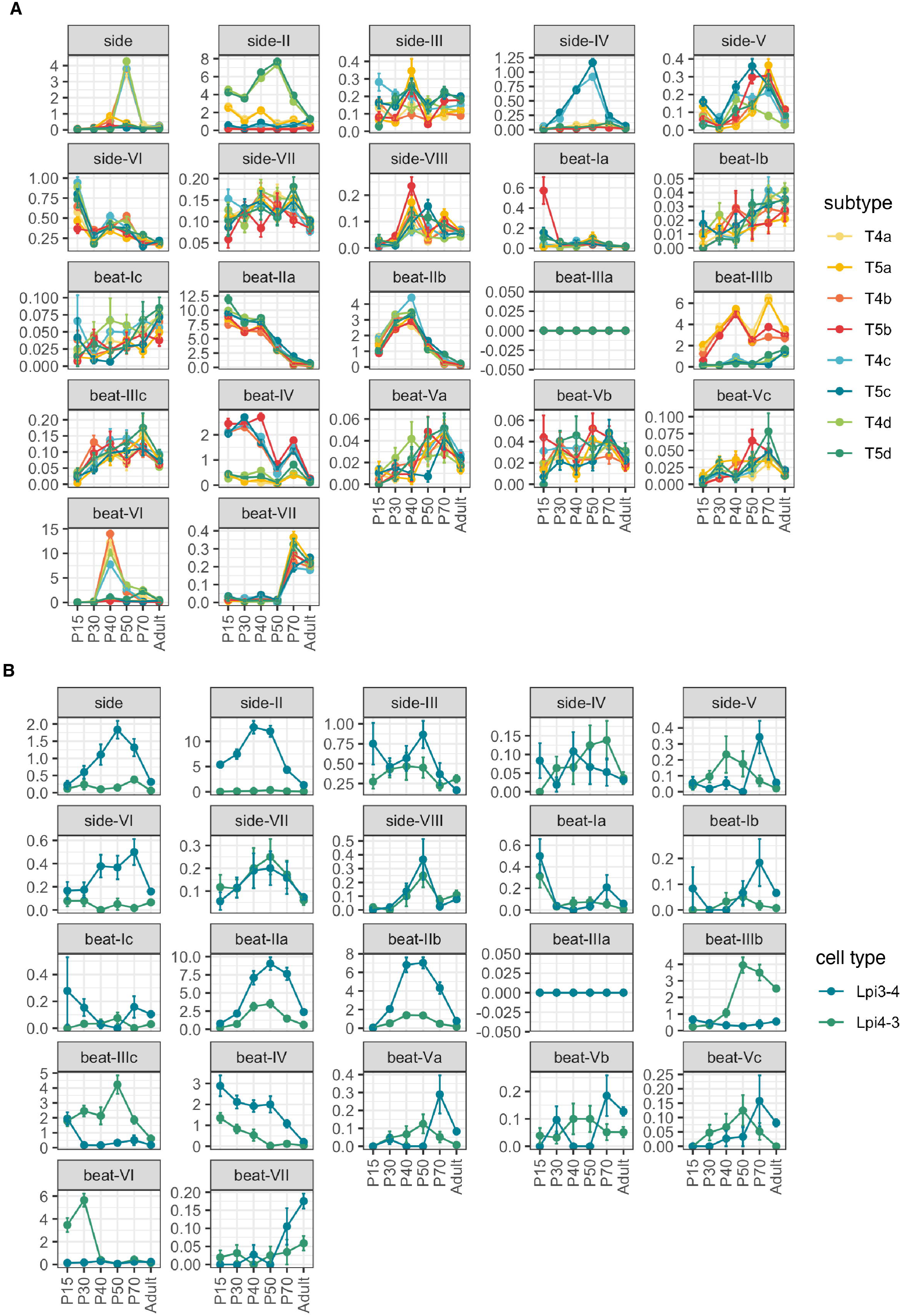
Beat/Side expression in T4/T5 and LPi3-4 and LPi4-3 neurons. Line plots with average expression levels of beat/side in the T4/T5s (**A**), and LPi3-4 and LPi4-3 neurons (**B**), from the (Ozel et al., 2021) dataset.

**Figure S7.**
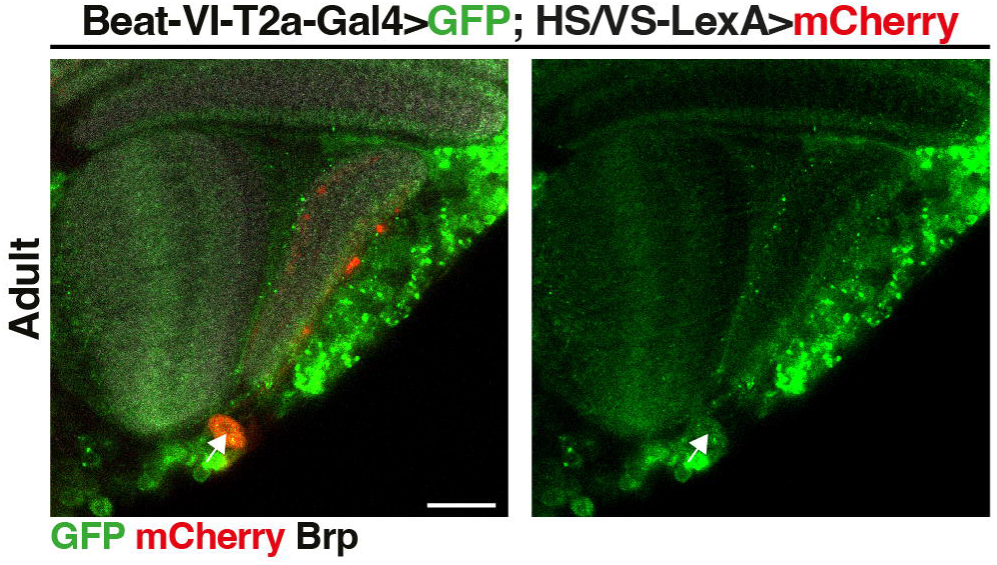
VS neurons express beat-VI. Beat-VI-T2A-Gal4 expressing cytosolic 6XGFP and HS/VS neurons labeled with HS/VS-LexA (R39E01-LexA) driving mcherry expression. VS neurons cell bodies colocalize with Beat-VI-T2A-Gal4 derived GFP expression. Scale bar is 20μm

**Figure S8.**
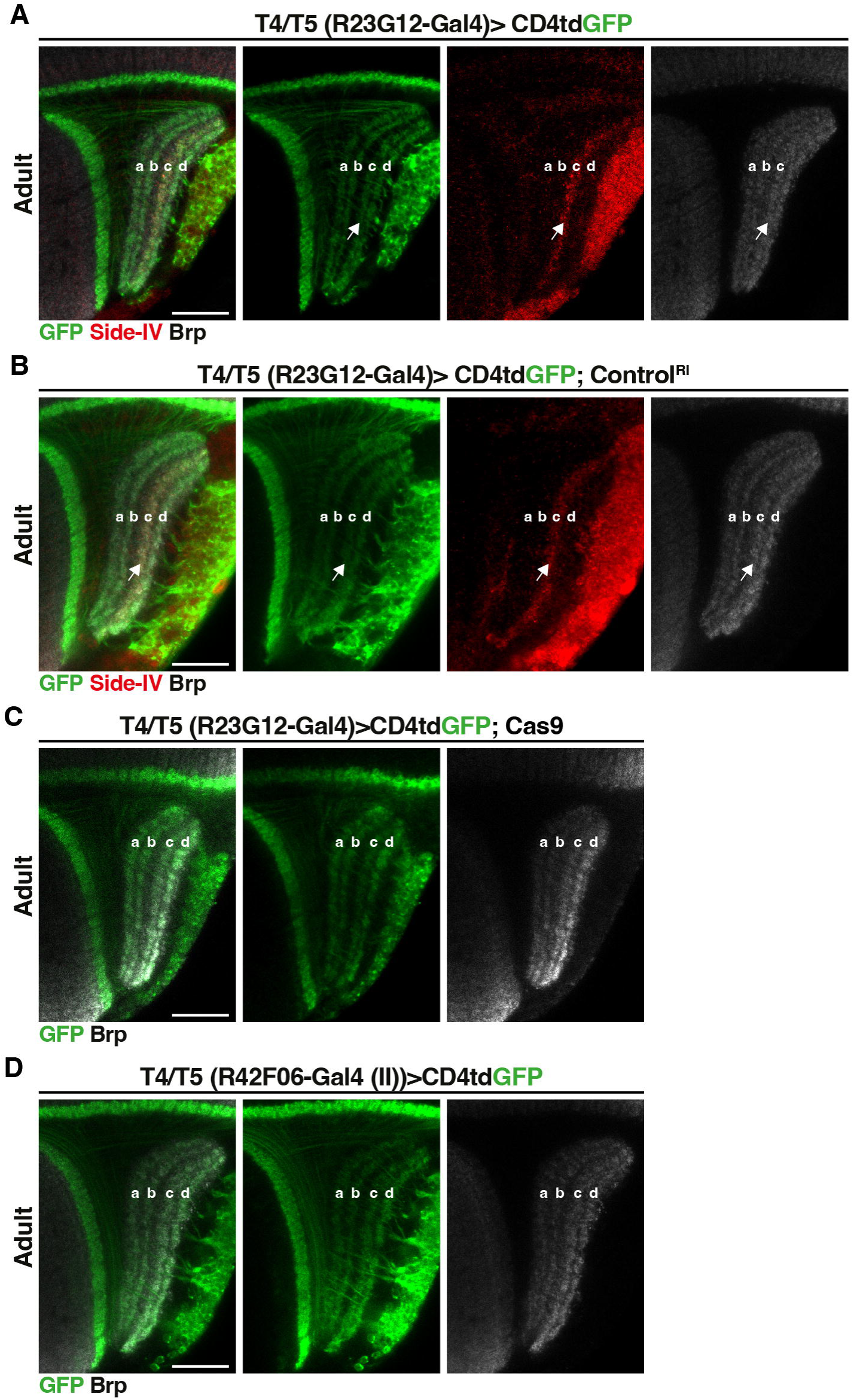
T4/T5-Gal4 Control lines. T4/T5 control lines for images 4 and 5 driving the expression of membrane tethered CD4tdGFP (green). Brp (grey) labels the neuropiles. In (**A**) and (**B**) Immunostaining against Side-IV (red) (arrow). (**A**) T4/T5 (R23G12-Gal). (**B**) T4/T5 (R23G12-Gal) driving control RNAi for side-II, side-IV and beat-VI RNAi experiments. (**C**) T4/T5 (R23G12-Gal) driving cas9 expression, control for sgRNA experiments. (**D**) T4/T5 (R42F06-Gal on the 2^nd^ chromossome), control for side-IV^null^; side-VI^null^; side-IV^null^, side-VI^null^; side-IV^null^, and beat-VI^null^ homozygous mutants conditions. Scale bars are 20μm

**Figure S9.**
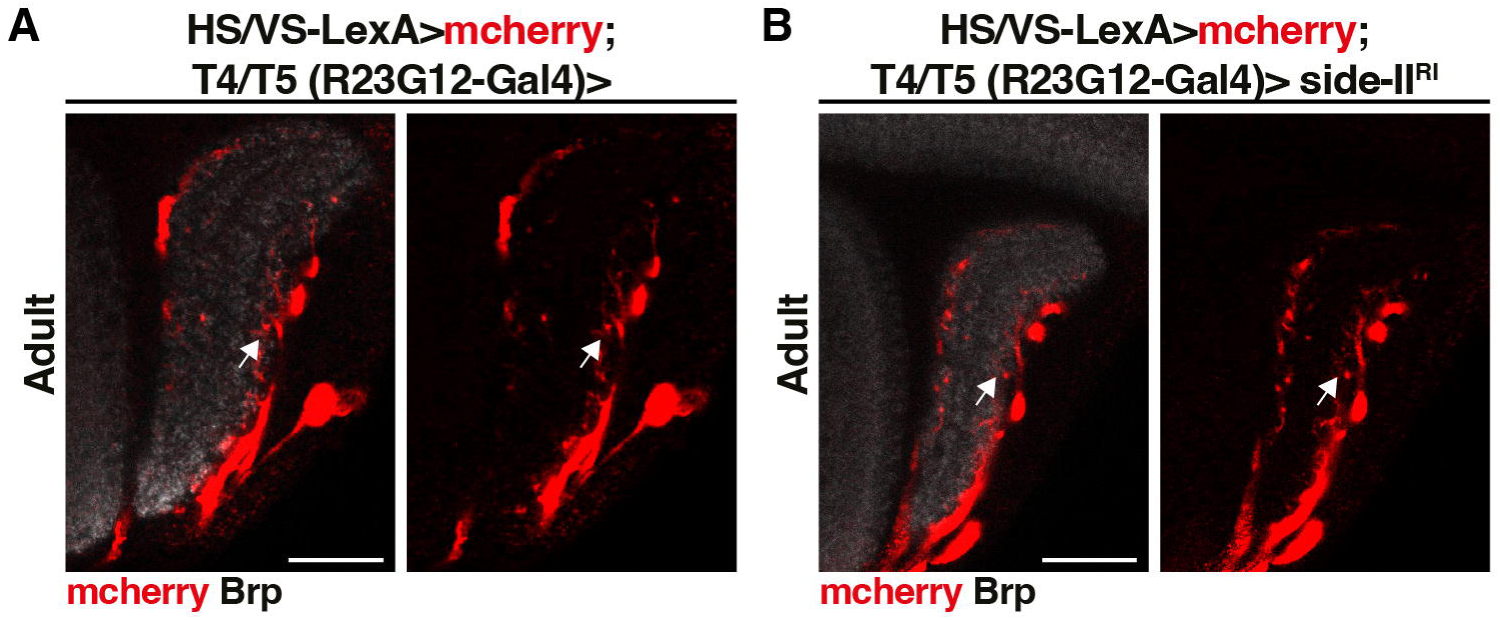
Side-II downregulation in T4/T5 neurons does not affect VS neurons layer innervation. HS/VS visualized with HS/VS-LexA (R39E01-LexA) driving mcherry. HS/VS neurons in the wildtype (**A**) and upon Side-II-RNAi mediated downregulation in T4/T5s using R23G12-Gal4 (**B**). Side-II downregulation does not affect HS/VS neurons layer innervation. Brp (grey) labels the neuropiles. Scale bars are 20μm.

**Figure S10.**
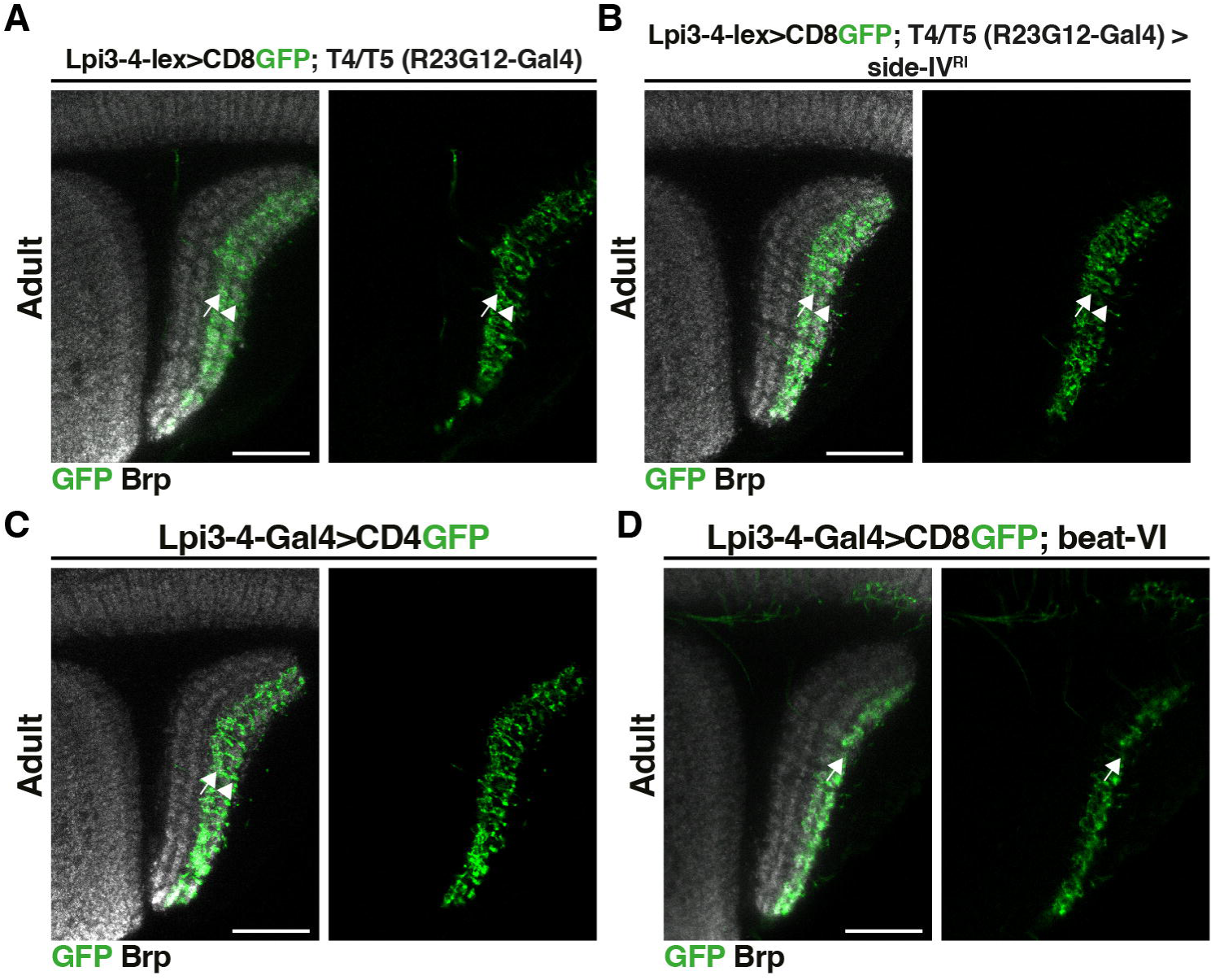
Beat-VI overexpression in LPi3-4 neurons is sufficient to change layer specific innervation. (**A**, **B**) Downregulation of Side-IV with RNAi in T4/T5 neurons does not affect LPi3-4 layer innervation. (**C**, **D**) Beat-VI overexpression in LPi3-4 promotes single layer 4 targeting. Arrow points to LPi3-4 dendritic processes, arrowhead to axons. Brp labels the neuropiles. Scale bars are 20μm.

