## Supplementary material for "Biased cell adhesion organizes a circuit for visual motion integration": Resource Table

| | | | | |
|---|---|---|---|---|
| **REAGENT or RESOURCE** | **SOURCE** | **IDENTIFIER** | **RRID full citation info** | **ADDITIONAL INFORMATION** |
| **Primary Antibodies** | | | | |
| rat anti-Side-IV | | | | Gift C. Desplan |
| guinea pig anti-Brp | | | | Gift C. Desplan |
| mouse anti-nc82 | | RRID:AB_2314866 | (DSHB Cat# nc82, RRID:AB_2314866) | |
| mouse anti-Connectin | DSHB | AB_10660830 | (DSHB Cat# Connectin C1.427, RRID:AB_10660830) | |
| rabbit anti-GFP | Biorad | AB_2313770 | (Torrey Pines Biolabs Cat# TP401, RRID:AB_2313770) | |
| chicken anti-GFP | Abcam | AB_300798 | (Abcam Cat# ab13970, RRID:AB_300798) | |
| sheep anti-GFP | Biorad | AB_619712 | (Bio-Rad Cat# 4745-1051, RRID:AB_619712) | |
| Phalloidin 405 | | Invitrogen™ A30104 | Invitrogen™ A30104 | |
| rabbit anti-RFP | tebu-bio | 600-401-379 | (Rockland Cat# 600-401-379, RRID:AB_2209751) | |
| chicken anti-V5 | Abcam | AB_307022 | (Abcam Cat# ab9113, RRID:AB_307022) | |
| rat anti-DNcad | DSHB | AB_528121 | (DSHB Cat# DN-Ex #8, RRID:AB_528121) | |
| **Secondary Antibodies** | | | | |
| see Table 2 | | | | |
| **Single-cell transcriptional atlas** | | | | |
| Single-cell transcriptional atlas of the Drosophila visual system | Ozel et al.; 2021 | GSE142787 | GSE142787 | |
| Single-cell transcriptional atlas of the Drosophila visual system | Yoo. Et al. 2023 | GSE156455 | GSE156455 | |
| **Plasmids** | | | | |
| FlpTag plasmids | Fendl et al.; 2020 | | | Gift A. Borst |
| **Experimental models: Organisms/strains** | | | | |
| Fly: D. melanogaster: T4/T5-Gal4 (R42F06) | BDSC | BDSC #41253 | BDSC_41253 | |
| Fly: D. melanogaster: T4/T5-Gal4 (R23G12) | BDSC | BDSC #49044 | BDSC _49044 | |
| Fly: D. melanogaster: T4/T5ad-Gal4 | This study | N/A | | Gift A. Borst |
| Fly: D. melanogaster: TmY14-Gal4 | Chen et al.; 2023 | N/A | | Gift C. Desplan |
| Fly: D. melanogaster: Y3-like-Gal4 | Chen et al.; 2023 | N/A | | Gift C. Desplan |
| Fly: D. melanogaster: LLPC2/LPC2-Gal4 (R80G09) | This study | BDSC #40089 | BDSC_40089 | |
| Fly: D. melanogaster: LPi3-4-Split-Gal | This study | | | |
| Fly: D. melanogaster: LPi3-4-LexA (R20D01) | Mauss et al.; 2015 | BDSC #61518 | BDSC_61518 | |
| Fly: D. melanogaster: LPi4-3-Gal4 (R38G02) | Mauss et al.; 2015 | BDSC #50016 | BDSC_50016 | |
| Fly: D. melanogaster: LPi4-3-Split-Gal4 (SS03752) | Klapoetke et al., 2017 | BDSC #76004 | BDSC_76004 | |

| | | | | |
|---|---|---|---|---|
| Fly: D. melanogaster: LPi4-3-LexA (R38G02) | Mauss et al.; 2015 | BDSC #52772 | BDSC_52772 | |
| Fly: D. melanogaster: HS/VS-Gal4 (R39E01) | Fujiwara et al.; 2017 | BDSC #50048 | BDSC_50048 | Gift E. Chiappe |
| Fly: D. melanogaster: HS/VS-LexA (R39E01) | Fujiwara et al.; 2017 | BDSC #52776 | BDSC_52776 | Gift E. Chiappe |
| Fly: D. melanogaster: hsFlp,Tubgal80 Frt19a;; | | | | Gift C. Desplan |
| Fly: D. melanogaster: hsFlp;Tubgal80 Frt40; | | | | Gift C. Desplan |
| Fly: D. melanogaster: Frt19a;; | BDSC | BDSC #1709 | BDSC_1709 | Gift C. Desplan |
| Fly: D. melanogaster: ;Frt40; | | BDSC #5192 | BDSC_5192 | |
| Fly: D. melanogaster: ;uas-cd4tdGFP; | | | | Gift C. Desplan |
| Fly: D. melanogaster: ;;uas-cd4tdGFP | | | | Gift C. Desplan |
| Fly: D. melanogaster: ;;uas-cd4tdTOMATO | | | | Gift C. Desplan |
| Fly: D. melanogaster: 10XUAS-IVS-mCD8::RFP, 13XLexAop2-mCD8::GFP;; | BDSC | BDSC #32229 | BDSC_32229 | |
| Fly: D. melanogaster: 10XUAS-IVS-mCD8::RFP, 13XLexAop2-mCD8::GFP | BDSC | BDSC #32229 | BDSC_32229 | |
| Fly: D. melanogaster:13XLexAop2-6XmCherry-HA;; | BDSC | BDSC #52272 | BDSC_52272 | Gift E. Chiappe |
| Fly: D. melanogaster::;10XUAS-IVS-myr::tdTomato | BDSC | BDSC #32221 | BDSC_32221 | |
| Fly: D. melanogaster: ;;UAS-Da7::GFP | Raghu et al., 2009 | | | Gift A. Borst |
| Fly: D. melanogaster: ;UAS-DenMark, UAS-syt.eGFP; | BDSC | BDSC #33064 | BDSC_33064 | |
| Fly: D. melanogaster: ;side-IInull FRT40; | This study | | | |
| Fly: D. melanogaster: ;uas-side-IV::V5 | Osaka et al.; 2023 | N/A | N/A | Gift T. Suzuki |
| Fly: D. melanogaster: ;;side-IVnull | Osaka et al.; 2023 | N/A | N/A | Gift T. Suzuki |
| Fly: D. melanogaster: ;;side-VIInull | Osaka et al.; 2023 | N/A | N/A | Gift T. Suzuki |
| Fly: D. melanogaster: ;;side-IVnull, side-VIInull | Osaka et al.; 2023 | N/A | N/A | Gift T. Suzuki |
| Fly: D. melanogaster: ;;beat-VInull | Osaka et al.; 2023 | N/A | N/A | Gift T. Suzuki |
| Fly: D. melanogaster: ;uas-side-VII; | Osaka et al.; 2023 | N/A | N/A | Gift T. Suzuki |
| Fly: D. melanogaster: ;uas-beat-IIa; | Osaka et al.; 2023 | N/A | N/A | Gift T. Suzuki |
| Fly: D. melanogaster: ;;beat-IIa::FlpTag-Constitutive | This study | | | |
| Fly: D. melanogaster: ;side-II::FlpTag-Constitutive; | This study | | | |
| Fly: D. melanogaster: ;;beat-IIa MIMIC | BDSC | BDSC #24550 | RRID:BDSC_24550 | |
| Fly: D. melanogaster: ;side-II MIMIC; | BDSC | BDSC #43603 | RRID:BDSC_43603 | |
| Fly: D. melanogaster: ;;beat-IV::GFSTF | BDSC | BDSC #66506 | BDSC_66506 | |
| Fly: D. melanogaster: ;;UAS-Dcr-2.D | BDSC | BDSC #24651 | BDSC_24651 | |
| Fly: D. melanogaster: ;;beat-VI-T2A-Gal4 | | | | Gift R. Mann |
| Fly: D. melanogaster: UAS-RNAi of side-IV: GD5643 | VDRC | VDRC#GD-16636 | VDRC#GD-16636 | |
| Fly: D. melanogaster: UAS-RNAi of side-IV: KK111999 | VDRC | VDRC#KK-102563 | VDRC#KK-102563 | |
| Fly: D. melanogaster: UAS-RNAi of side-II: KK111520 | VDRC | VDRC#KK-107512 | VDRC#KK-107512 | |

| | | | |
|---|---|---|---|
| Fly: D. melanogaster: UAS-RNAi of side-II: KK114253 | VDRC | KK-103687 | KK-103687 |
| Fly: D. melanogaster: UAS-RNAi of beat-VI: GD89 | VDRC | VDRC#GD-6694 | VDRC#GD-6694 |
| Fly: D. melanogaster: UAS-RNAi of beat-VI: KK111999 | VDRC | KK-102563 | KK-102563 |
| Fly: D. melanogaster: empty attP control line | VDRC | VDRC#TK-60100 | VDRC#TK-60100 |
| Fly: D. melanogaster: MCFO-1 | BDSC | BDSC #64085 | BDSC_64085 |
| Fly: D. melanogaster: UAS-Stinger, UAS-hid.Z | BDSC | BDSC #65408 | BDSC_65408 |
| Fly: D. melanogaster: UAS-beat-VI.ORF.3xHA | FlyORF | F002906 | F002906 |

Table-II Secondary antibodies used

| Fluorophore | Host | Target | name | supplier / ref | RRID |
|---|---|---|---|---|---|
| Alexa Fluor 488 | Donkey | anti-Rat | ALEXA FLUOR 488 DONKEY ANTI RAT IgG (H+L) | Invitrogen A21208 | (Thermo Fisher Scientific Cat# A-21208, RRID:AB_2535794) |
| Alexa Fluor 488 | Donkey | anti-Goat | ALEXA FLUOR 488 DONKEY ANTIGOAT IgG (H+L) | Invitrogen A11055 | (Thermo Fisher Scientific Cat# A-11055 (also A11055), RRID:AB_2534102) |
| Alexa Fluor 555 | Donkey | anti-Goat | ALEXA FLUOR 555 DONKEY ANTI GOAT IgG (H+L) | Invitrogen A21432 | (Thermo Fisher Scientific Cat# A-21432 (also A21432), RRID:AB_2535853) |
| Alexa Fluor 555 | Donkey | anti-Mouse | ALEXA FLUOR 555 DONKEY ANTI MOUSE IgG (H+L) | Invitrogen A31570 | (Thermo Fisher Scientific Cat# A-31570, RRID:AB_2536180) |
| Alexa Fluor 488 | Donkey | anti-Rabbit | ALEXA FLUOR 488 DONKEY ANTI RABBIT IgG (H+L) | Invitrogen A21206 | (Thermo Fisher Scientific Cat# A-21206 (also A21206), RRID:AB_2535792) |
| Alexa Fluor 488 | Donkey | anti-Sheep | ALEXA FLUOR 488 DONKEY ANTI SHEEP IgG (H+L) | Invitrogen A11015 | (Thermo Fisher Scientific Cat# A-11015, RRID:AB_2534082) |
| Alexa Fluor 647 | Donkey | anti-Goat | ALEXA FLUOR 647 DONKEY ANTI GOAT IgG (H+L) | Invitrogen A21447 | (Thermo Fisher Scientific Cat# A-21447, RRID:AB_2535864) |
| Alexa Fluor 647 | Donkey | anti-Rabbit | ALEXA FLUOR 647DONKEY ANTI RABBIT IgG (H+L) | Invitrogen A31573 | (Thermo Fisher Scientific Cat# A-31573, RRID:AB_2536183) |
| Alexa Fluor 488 | Donkey | anti-Mouse | ALEXA FLUOR 488 DONKEY ANTI MOUSE IgG (H+L) | Invitrogen A21202 | (Thermo Fisher Scientific Cat# A-21202, RRID:AB_141607) |
| Alexa Fluor 555 | Donkey | anti-Sheep | ALEXA FLUOR555 DONKEY ANTI SHEEP IgG (H+L) | Invitrogen A21436 | (Thermo Fisher Scientific Cat# A-21436, RRID:AB_2535857) |
| Alexa Fluor 555 | Donkey | anti-Rabbit | ALEXA FLUOR555 DONKEY ANTI RABBIT IgG (H+L) | Invitrogen A31572 | (Thermo Fisher Scientific Cat# A-31572 (also A31572), RRID:AB_162543) |
| Rhodamine red X-570nm | Donkey | anti-Chicken | Rhodamine Red™-X (RRX) AffiniPure Donkey Anti-Chicken IgY (IgG) (H+L) | Jackson ImmunoResearch 703-295-155 | (Jackson ImmunoResearch Labs Cat# 703-295-155, RRID:AB_2340371) |
| Alexa Fluor 488 | Donkey | anti-Chicken | ALEXA FLUOR 488 AFFINIPURE DONKEY ANTI CHICKEN IgY | Jackson ImmunoResearch 703-545-155 | (Jackson ImmunoResearch Labs Cat# 703-545-155, RRID:AB_2340375) |
| Alexa Fluor 647 | Donkey | anti-Chicken | ALEXA FLUOR 647 AFFINIPREDONKEY ANTI CHICKEN IgY | Jackson ImmunoResearch 703-605-155 | (Jackson ImmunoResearch Labs Cat# 703-605-155, RRID:AB_2340379) |
| Cy3-532nm | Donkey | anti-Guinea pig | CY3 AFFINIPURE DONKEY ANTI GUINEA PIG IgG | Jackson ImmunoResearch 706-165-148 | (Jackson ImmunoResearch Labs Cat# 706-165-148, RRID:AB_2340460) |
| Alexa Fluor 488 | Donkey | anti-Guinea pig | ALEXA FLOUR 488-AFFINIPRE DONKEY ANTI GUINEA PIG IgG | Jackson ImmunoResearch 703-545-155 | (Jackson ImmunoResearch Labs Cat# 703-545-155, RRID:AB_2340375) |
| Alexa Fluor 647 | Donkey | anti-Guinea pig | Alexa Fluor® 647 AffiniPure Donkey Anti-Guinea Pig IgG (H+L | Jackson ImmunoResearch 706-605-148 | (Jackson ImmunoResearch Labs Cat# 706-605-148, RRID:AB_2340476) |
| Cy3-532nm | Donkey | anti-Rat | CY3 AFFINIPUREDONKEY ANTI RAT IgG | Jackson ImmunoResearch 712-165-153 | (Jackson ImmunoResearch Labs Cat# 712-165-153, RRID:AB_2340667) |
| DyeLight 405 | Donkey | anti-Rat | DYLIGHT AFFINI PURE DONKEY ANTIRAT IgG 0.5 mg | Jackson ImmunoResearch 712-475-150 | (Jackson ImmunoResearch Labs Cat# 712-475-150, RRID:AB_2340680) |

| | | | | | |
|---|---|---|---|---|---|
| Alexa Fluor 647 | Donkey | anti-Rat | ALEXA FLOUR 647 AFFINIPURE F(AB)2 FRAGMENT DONKEY ANTI RAT IgG | Jackson ImmunoResearch 712-606-150 | (Jackson ImmunoResearch Labs Cat# 712-606-150, RRID:AB_2340695) |
| DyeLight 405 | Donkey | anti-Mouse | DyLight™ 405 AffiniPure Donkey Anti-Mouse IgG (H+L) | Jackson ImmunoResearch 715-475-150 | (Jackson ImmunoResearch Labs Cat# 715-475-150, RRID:AB_2340839) |
| Alexa Fluor 647 | Donkey | anti-Mouse | ALEXA FLUOR 647 DONKEY ANTI MOUSE IgG (H+L) | Jackson ImmunoResearch 715-605-140 | (Jackson ImmunoResearch Labs Cat# 715-605-140, RRID:AB_2340861) |
| Alexa Fluor 488 | Goat | Anti-Rabbit | Highly Cross-Adsorbed Goat anti-Rabbit, Alexa Fluor™ 488 | Invitrogen A11034 | (Thermo Fisher Scientific Cat# A-11034 (also A11034), RRID:AB_2576217) |
| Alexa Fluor 555 | Goat | Anti-Rabbit | Highly Cross-Adsorbed Goat anti-Rabbit, Alexa Fluor™ 555 | Invitrogen A21428 | (Thermo Fisher Scientific Cat# A-21428, RRID:AB_2535849) |
| Alexa Fluor 647 | Goat | Anti-Rabbit | Highly Cross-Adsorbed Goat anti-Rabbit, Alexa Fluor™ 647 | Invitrogen A21244 | (Thermo Fisher Scientific Cat# A-21244, RRID:AB_2535812) |
| Dylight 405 | Donkey | Anti-Rabbit | DyLight 405-AffiniPure Donkey Anti-Rabbit IgG (H+L) | Jackson ImmunoResearch 711-475-152 | (Jackson ImmunoResearch Labs Cat# 711-475-152, RRID:AB_2340616) |
| Alexa Fluor 647 | Goat | Anti-Mouse | IgG (H+L) Cross-Adsorbed Goat anti-Mouse, Alexa Fluor™ 647, Invitrogen | Invitrogen A21235 | (Thermo Fisher Scientific Cat# A-21235, RRID:AB_2535804) |
| Alexa Fluor Plus 555 | Donkey | Anti-Mouse | IgG (H+L) Highly Cross-Adsorbed Donkey anti-Mouse, Alexa Fluor™ Plus 555, | Invitrogen A32773 | (Thermo Fisher Scientific Cat# A32773, RRID:AB_2762848) |
| Alexa Fluor Plus 647 | Donkey | Anti-Mouse | IgG (H+L) Highly Cross-Adsorbed Donkey anti-Mouse, Alexa Fluor™ Plus 647, | Invitrogen A32787 | (Thermo Fisher Scientific Cat# A32787, RRID:AB_2762830) |
